# Overexpression of α-Synuclein Alters The Nanoscopic Organisation of Presynaptic Proteins

**DOI:** 10.1101/2025.10.10.681706

**Authors:** Iain A. Watson, Jessika C. Bridi, Diane Hanger, Frank Hirth, Deepak P. Srivastava

## Abstract

Parkinson’s Disease (PD) is characterised by accumulation of α-synuclein (α-Syn), but how elevated α-Syn alters presynaptic architecture during prodromal phases of the disease remains unclear. We investigated presynaptic morphology and two key presynaptic proteins: MYCBP2; and the Active Zone (AZ) protein ELKS. Primary rat cortical neurons were transfected with wildtype α-Syn, or the familial A30P mutant, and nanoscopic changes to bouton morphology and protein localisation were investigated using super-resolution microscopy. Variant specific accumulation patterns for overexpressed α-Syn were observed without changes to bouton structure. Additionally, increases of both α-Syn variants affected presynaptic proteins, decreasing MYCBP2 puncta count and intensity, and reducing ELKS protein density. Since MYCBP2 potentially regulates ELKS, these findings suggest that elevated α-Syn perturbs AZ components whilst presynaptic structure is preserved. Our study supports growing evidence that increased α-Syn disturbs presynaptic function potentially through a MYCBP2-ELKS axis, a mechanism which may provide a valuable target for PD therapeutics.

## Introduction

The α-synuclein (α-Syn)-encoding *SNCA* gene has consistently emerged as a significant genetic risk factor for Parkinson’s disease (PD) in both familial and genome-wide association studies (Aarsland et al., 2005; Chang et al., 2017; Golbe et al., 1990; Jia et al., 2022; Nalls et al., 2014, 2019; Trinh et al., 2018). Specific point mutations in *SNCA*, including A30P, A53T, and E46K, have been linked to familial and early-onset forms of PD (Appel-Cresswell et al., 2013; Krüger et al., 1998; Lesage et al., 2013; Pasanen et al., 2014; Polymeropoulos et al., 1997; Zarranz et al., 2004). At a pathological level, α-Syn aggregation is a defining pathological hallmark of PD, leading to formation of Lewy bodies within affected neurons (Braak et al., 1999; Goedert & Spillantini, 1998; Polymeropoulos et al., 1997; Spillantini et al., 1997). Beyond its pathological implications, α-Syn plays important physiological roles in regulating synaptic vesicle trafficking, neurotransmitter release, and presynaptic organisation (Cabin et al., 2002; Fouke et al., 2021; Hoffmann et al., 2021). Under physiological conditions, α-Syn localises to presynaptic sites (Iwai et al., 1995), where it contributes to synaptic homeostasis. In PD and related synucleinopathies, however, α-Syn becomes abnormally accumulated at these presynaptic sites, potentially disrupting vesicle dynamics and neurotransmission (Kramer & Schulz-Schaeffer, 2007). Despite these insights, the precise mechanisms by which pathological aggregation of α-Syn alters presynaptic structure or function in PD remains poorly understood. Elucidating these mechanisms is essential for understanding how α-Syn contributes to PD pathogenesis.

At the presynapse, synaptic boutons come together to form specialised sites dedicated to communication with postsynaptic cells. The active zone (AZ) within presynaptic boutons plays a pivotal role in regulating neurotransmission and synaptic vesicle exocytosis. Comprising a dense network of evolutionarily conserved proteins, the AZ serves as a molecular scaffold that organises and coordinates the processes underlying synaptic transmission. Among these proteins, ELKS emerges as a key component, functioning as a scaffold protein anchoring and stabilising other AZ proteins to maintain normal synaptic function (Ohtsuka, 2013).

In a *Drosophila* model of synucleinopathy, α-Syn overexpression lead to its accumulation at the neuromuscular junction (NMJ), where it altered the expression of Bruchpilot (BRP), a core component of the neuronal AZ (Bridi et al., 2021). BRP belongs to a superfamily of proteins known as Rab3-interacting molecule-binding proteins (RIM-BPs), which act as highly conserved scaffolding proteins mediating synaptic vesicle release (Petzoldt et al., 2020; Zang et al., 2013). α-Syn accumulation at presynaptic terminals reduced the expression of BRP at the AZ, and differentially regulated other presynaptic proteins, including Synapsin and Syntaxin, effects primarily attributed to BRP dysfunction (Bridi et al., 2021). In mammals, ELKS is the structural and functional homologue of BRP (Wagh et al., 2006). However, whether α-Syn can modulate the expression or localisation of ELKS within the mammalian presynaptic AZs remains unknown.

Previous research has shown that BRP can be regulated through a pathway in *Drosophila* involving Highwire and dNmnat (Russo et al., 2019). Highwire controls the size of the presynaptic terminal at the NMJ, without altering BRP puncta number (Russo et al., 2019). Other studies have shown that dNmnat is critical for maintaining the structural integrity of the AZ via BRP (Zang et al., 2013). Downregulation of the mammalian homologue to dNmnat, NMNAT, has been implicated in several neurodegenerative diseases, including PD, Alzheimer’s Disease (AD), and Huntington’s Disease (HD) (Ali et al., 2013). The mammalian homologue of Highwire is MYCBP2. But whether MYCBP2 can similarly regulate the AZ of mammalian neurons remains an open question. Together, these observations suggest that the Highwire–dNmnat pathway and its regulation of AZ components in *Drosophila* may represent a conserved mechanism potentially exploited by α-Syn pathology in synucleinopathies (Bridi & Hirth, 2018).

In this study, we sought to investigate whether elevated levels of α-Syn, modelling the heightened protein levels characteristic of Parkinsonian pathology through the overexpression of α-Syn in neuronal cultures (Jakes et al., 1994; Spillantini et al., 1997), influences the nanoscopic organisation of the mammalian presynaptic AZ. Although α-Syn has been extensively implicated in synaptic dysfunction (Bridi et al., 2021; Chandra et al., 2004; L. Chen et al., 2022; Greten-Harrison et al., 2010; Kramer & Schulz-Schaeffer, 2007; Pérez-Acuña et al., 2023; Vargas et al., 2017; Q. Wu et al., 2019), the mechanisms by which it affects presynaptic structure and signalling remain an active area for further examination (Bridi & Hirth, 2018; Y.-J. Chen et al., 2025; Song et al., 2023; Uytterhoeven et al., 2024; Vargas et al., 2025; Yoo et al., 2023). Evidence from *Drosophila* synucleinopathy models implicates a link between α-Syn and the dNmnat/Highwire signalling pathway; however, whether the mammalian counterpart, MYCBP2, is also disrupted following α-Syn accumulation is unclear. Moreover, it is unknown whether the accumulation of α-Syn or its mutant forms differentially affect the AZ or related protein organisation at the presynapse. To address these gaps, we examined whether overexpression of wildtype (WT) or mutant α-Syn alters the expression or localisation of MYCBP2 and the AZ scaffolding protein ELKS, thereby impacting the nanoscopic organisation and architecture of presynaptic boutons. We employed instant Structured Illumination Microscopy (iSIM) to visualise presynaptic boutons at high resolution, providing detailed insight into AZ organisation under conditions of α-Syn overexpression. Through super resolution imaging, we assessed structural and organisational changes within presynaptic terminals, focusing on alterations in AZ morphology and protein composition. This approach enabled us to define the relationship between α- Syn overexpression and key AZ components, providing new insights into how α-Syn may modulate presynaptic integrity and signalling in the mammalian central nervous system.

## Methods

### Reagents

Primary antibodies used were: GFP chicken polyclonal (ab13970; Abcam; 1:10000), MycBP2 rabbit polyclonal (ab86078; Abcam; 1:100), ELKS mouse monoclonal antibody (Clone ELKS-30; E4531; Sigma; 1:500), MAP2 chicken polyclonal (ab92434; Abcam; 1:500), Synapsin-1 rabbit monoclonal antibody (Clone D12G5; 5297; Cell Signaling; 1:1000), Bassoon mouse monoclonal antibody (Clone SAP7F407; ab82958; Abcam; 1:200). Secondary antibodies used: anti-Chicken Alexa Fluor 488 goat polyclonal (A-11034; Thermo; 1:1000); anti-Rabbit Alexa Fluor 568 goat polyclonal (A-11011; Thermo; 1:500); anti-Mouse Alexa Fluor 680 goat polyclonal (A-21058; Thermo; 1:500). Plasmids include the use of enhanced GFP-C3 (eGFP), α-synuclein-wildtype (α-Syn-WT) and mutant α-synuclein-A30P (α-Syn- A30P) kindly donated by Frank Hirth and Diane Hanger (Bridi et al., 2021; Paillusson et al., 2017). Plasmids were prepared using the ZymoPURE II Plasmid kit Maxiprep Kit (D4202; Zymo Research) following the manufacturers guidelines.

### Cell culture and transfections

Primary rat cortical neurons were prepared from E18 Sprague-Dawley embryos as previously described (Srivastava et al., 2011). Animals were habituated for 3 days prior to experimental procedures, and all procedures were carried in accordance with the Home Office Animals (Scientific Procedures) Act, United Kingdom, 1986. All animal experiments were given ethical approval by the ethics committee of King’s College London (United Kingdom). Neurons were then seeded as mixed sex cultures onto poly-D-lysine (0.2 mg/ml, Sigma) coated 18mm glass coverslips (No 1.5; 0117580, Marienfeld-Superior GmbH & Co.) at a density of 2.5x105/well, equivalent to 714/mm2. Neurons were cultured with the following media: neurobasal medium (21103049) supplemented with 2% B27 (17504044), 0.5 mM glutamine (25030024) and 1% penicillin:streptomycin (15070063) (all reagents from Life Technologies). Half media changes were made twice weekly until the neurons were at the required age of Days In Vitro (DIV) 23. Neurons were transfected using the required plasmids at DIV20 for 3 days using Lipofectamine 2000 (11668027, Life Technologies) following manufacturers guidelines. 1 µg of plasmid DNA was mixed with 2 µl Lipofectamine 2000 and incubated for 4-12 hours in antibiotic-free media, after transfection the cells were replaced in original media for a further 3 days before fixation.

### Immunocytochemistry

After transfection neurons were fixed in a 4% paraformaldehyde/4% sucrose Phosphate Buffered Solution (PBS) solution for 10 minutes. This was followed by a post-fixation at 4^0^C with Methanol (pre-chilled to -20^0^C). Neurons were then simultaneously permeabilised and blocked using a 2% NGS (5425A; New England Biolabs) and 0.1% Triton X-100 PBS solution for one hour at room temperature. All antibodies were diluted in a 2% NGS in PBS blocking solution; neurons were incubated with primary antibodies overnight at 4^0^C or with secondary antibodies for one hour at room temperature. Neurons were washed three times for 10 minutes after each antibody incubation. Coverslips were finally mounted using ProLong Gold Antifade reagent (Life Technologies).

### Microscopy

Super-resolution images were obtained using Nikon iSIM ECLIPSE Ti-E inverted microscope using 100x oil immersion objective (Nikon, N.A. 1.49). Sections of secondary or tertiary dendrite were imaged from 3-5 cells per condition across 3 experiments. Imaging parameters were maintained within each biological repeat. Images were obtained as stacks of 51 slices with step sizes of 0.12 µm and subsequently deconvolved by blind deconvolution method using the in-built iSIM-specific deconvolution module in Nikon NIS-Elements Advanced Research (v5.01.00, Nikon). Two-dimensional maximum projection reconstructions of images were generated using ImageJ (https://imagej.nih.gov/ij/). Technical repeats for each experiment were conducted by imaging 3-5 cells per condition. The data from these technical repeats were averaged to generate a mean value for each condition, which was considered one biological replicate. These biological replicates are reported as the ’N’ number in the study. A total of 3-5 biological replicates were performed.

### Analysis

Analysis was carried out using the Conda package and environment management system (Anaconda Navigator v1.9.12, https://anaconda.org/, Anaconda, Inc.). Statistical analysis was performed using Python3 (v3.7.13, https://www.python.org/, Python Software Foundation) using the Spyder IDE (v5.2.2, https://www.spyder-ide.org/,Spyder Website Contributors). Additional Python modules were installed that were outside the base Anaconda environment, and included scikit-posthocs (v0.7.0, https://pypi.org/project/scikit-posthocs/) and researchpy (v0.3.2, https://pypi.org/project/researchpy/).

For multiple condition comparisons, ANOVA was used. Normality checks were conducted using the Jarque-Bera test to assess skewness and kurtosis, while the Shapiro-Wilk test was performed for each condition. Data sets were also visualized with Q-Q and histogram plots to confirm patterns and potential outliers. When ANOVA indicated significance, post-hoc tests were conducted using Tukey’s HSD method to pinpoint differences between groups. If normality failed, a single pre-specified Grubbs screen (α=0.05; no iteration) was used to flag potential outliers; flagged values were inspected for acquisition/segmentation artefacts and only artefacts were excluded. After outlier removal, ANOVA was repeated. If QC failed, non-parametric tests were applied. The Kruskal-Wallis H test was used for non-parametric multiple comparisons, followed by Dunn’s test if significant differences were found.

Analysis scripts can be found online https://github.com/iainawatson/Python-Analysis. Graphing was carried out using matplotlib (v3.5.2,https://matplotlib.org/, Matplotlib development team) and seaborn (v0.11.2, https://seaborn.pydata.org/, Michael Waskom). Box plots were created with whiskers to indicate 1.5x Interquartile Range (IQR), with additional dashed lines to indicate mean. Scatter plots were overlaid and colour coded according to biological repeats for clearer data visualisation. For multiple comparisons of conditions across two independent variables, two-way ANOVA were used. Error bars for two-way ANOVA are standard deviation.

## Results

### **α**-Synuclein transfections in rat cortical neurons

To explore the potential mechanisms of α-Syn at the presynaptic bouton, we conducted experiments using rat cortical neurons transfected with either wildtype or mutant A30P α-Syn overexpressions. The overexpression of α-Syn in neuronal cultures has previously been used to model aggregation of the protein characteristic of Parkinsonian pathology (Jakes et al., 1994; Spillantini et al., 1997). We optimised the amount of plasmid DNA to be used before testing the feasibility of identifying synaptic boutons in our neuronal cultures (Figure S1A). Under epifluorescent microscopy, we observed that overexpression of either eGFP or α-Syn-A30P did not produce obvious gross morphological effects of transfected neurons (Figure S1B, first column). Within these images, we could discern the axons and axon initial segments (indicated by white arrows in the first column of Figure S1B). However, due to the small size of these bouton structures, we used iSIM imaging to visualise synaptic boutons. We have previously shown that our iSIM microscope provides a lateral resolution of <160 nm and axial resolution of <370 nm (Gatford et al., 2021). With this form of super resolution microscopy, presynaptic boutons became distinctly visible along the length of the axon, consistently spaced at semi-regular intervals (Figure S1B, second column). These synaptic boutons closely associated with MAP2-positive stained neurons, indicating their potential role as sites for synaptic connections between different cells.

### ELKS is presynaptic marker of the Active Zone and associates with Synapsin-1

ELKS, also known as ERC1 or CAST, represents the vertebrate counterpart of the *Drosophila* protein BRP, known for its role in the localisation of AZ components such as RIM and Bassoon (Wagh et al., 2006). In previous studies, BRP was found to undergo alterations due to the accumulation of α-Syn within the NMJ (Bridi et al., 2021). In this context, our initial exploration focused on the visualisation of ELKS within presynaptic boutons. To achieve this, we coimmunostained rat cortical neurons with MAP2 to provide insight into the localisation of ELKS with the dendritic structure of the neurons. In addition, we utilised Synapsin-1, a well-characterised presynaptic marker within primary cortical neurons, to identify the presence of ELKS at the presynapse (Thiel, 1993). Synapsin-1 is closely associated with synaptic vesicle membranes and offers an effective means of visualising the presynaptic environment.

The images obtained through iSIM imaging were subjected to z-stacking and pseudo-coloured for enhanced clarity. ELKS immunostaining manifested as punctate signals distributed along dendritic regions (Figure 1A, first panel). In parallel, Synapsin-1 immunoreactivity displayed a pronounced punctate pattern along MAP2- postive dendrites (Figure 1A, second panel). Notably, at the sites where Synapsin-1 puncta were concentrated, ELKS staining frequently exhibited colocalisation, visually represented by white signals in the green and magenta colour scheme (Figure 1A, hollow arrows). While the z-stacked images effectively showcased the colocalisation between ELKS and Synapsin-1, this method might potentially misrepresent the depth of immunostaining acquisition. To substantiate the colocalisation of presynaptic content, orthogonal views were captured within regions suspected of colocalisation. This multi-angle perspective revealed that the colocalisation of ELKS and Synapsin- 1 was not limited to the xy axis but extended into the z axis of our images (Figure 1B, hollow arrows). This comprehensive observation establishes ELKS as a suitable antibody for our assay. It highlights the close association and colocalisation of both presynaptic markers, Synapsin-1 and ELKS, in the presynaptic regions surrounding dendrites.

**Figure 1.**
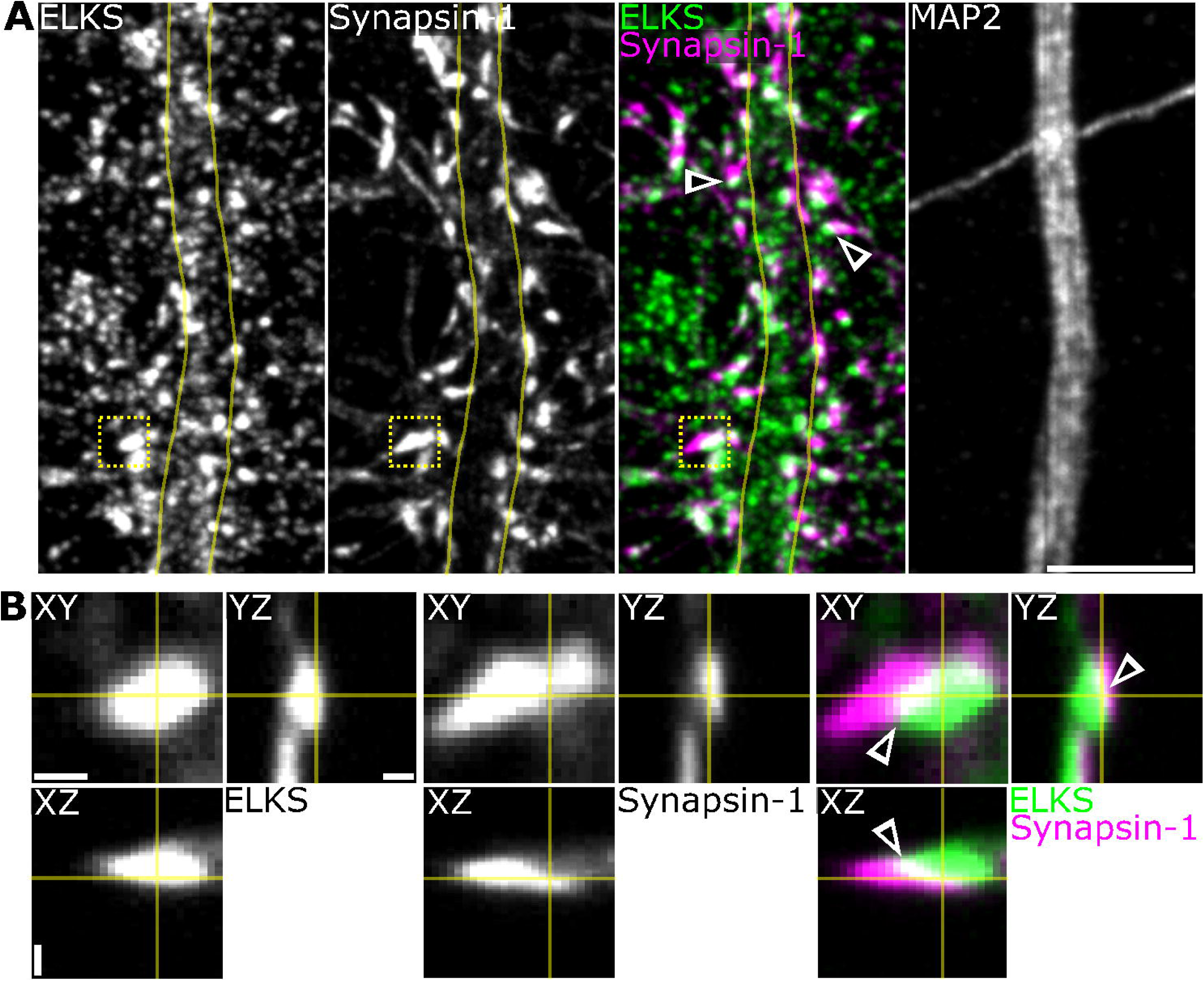
ELKS protein distribution. **(A)** Z-stacked images show sections of primary rat cortical neurons that were stained with the AZ marker ELKS. Cells were also immunostained with Synapsin-1 to confirm the presynapse and MAP2 to localise staining along sections of dendrite. In the colour merged images shades of white indicate colocalisation between ELKS and Synapsin-1, highlighted by arrows. **(B)** Orthogonal views are enlarged regions taken from the dashed yellow boxes indicated in (A) to show depth of the acquired stacks. The region of colocalisation is also given by arrows. Scale bar: (A) 5 µm; (B) 0.5 µm in XY, 1 µm in Z. Images acquired on iSIM.

### MYCBP2 is enriched presynaptically and associates with Bassoon

MYCBP2, a E3 ubiquitin-protein ligase, serves as the mammalian counterpart of the *Drosophila* protein Highwire. In *Drosophila*, dNmnat plays a pivotal role in regulating the ubiquitin-mediated degradation of the AZ protein BRP (Zang et al., 2013). Notably, the activity of dNmnat is regulated through a mechanism involving Highwire, which promotes the degradation of dNmnat (Xiong et al., 2012). This intricate process involving the Ubiquitin-Proteasome System potentially establishes link between the accumulation of α-Syn at the synapse and subsequent alterations to the AZ. To verify the localisation of MYCBP2 and its’ suitability for immunostaining, we co-stained MYCBP2 with Bassoon. Bassoon is a well-established presynaptic marker that exhibits robust staining in rat cortical neurons. Additionally, MAP2 was again used to delineate the dendritic structures in our neurons, providing contextual information for the observed staining patterns.

In our immunostaining observations, images were captured using iSIM and then subjected to z-projection. MYCBP2 exhibited abundant punctate staining pattern. Like ELKS, MYCBP2 displayed denser immunostaining along MAP2-positive dendrites when compared to the regions in between. This preferential proximity of MYCBP2 to MAP2 suggests its potential involvement in synaptic localisation (Figure 2). Upon overlaying pseudo-coloured images of MYCBP2 and Bassoon in green and magenta, it became evident that the antibody staining either closely associated or colocalised, indicated by the white colouration (Figure 2A, hollow arrows). For a more comprehensive view of this colocalisation in three dimensions, orthogonal views were generated (Figure 2B). The overlap of MYCBP2 and Bassoon was consistently retained within the z-axis (Figure 2B, hollow arrows). The punctate MYCBP2 staining appeared to localise around the Bassoon puncta. This level of clarity and the close association with Bassoon reaffirm that, within the parameters of our experiment, MYCBP2 is well-suited for our intended assay.

**Figure 2.**
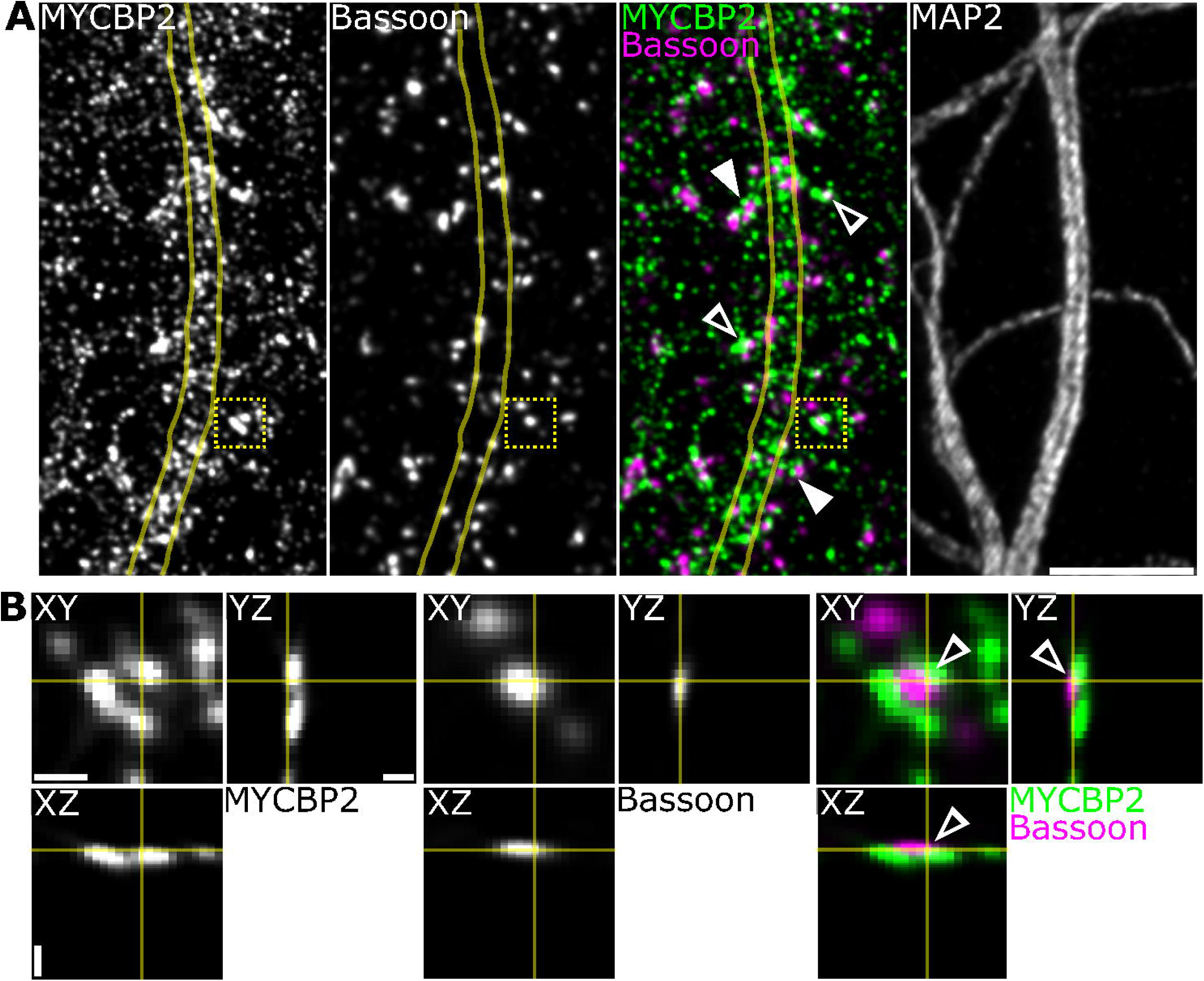
Distribution of MYCBP2 with the presynaptic marker Bassoon. **(A)** Z-stack images of rat cortical neurons immunostained for MYCBP2. MYCBP2 was coimmunostained with the presynaptic marker Bassoon to help identify presynaptic localisation. Cells were also coimmunostained with MAP2 to highlight areas of dendrite. In the merged image MYCBP2 can frequently be observed to associate closely with Bassoon as indicated by the solid arrows. Additionally, several areas are observed to colocalise as white in the merged image indicated by the hollow arrows. **(B)** Orthogonal views of the inset dashed yellow boxes in (A) to show the depth of the acquired images. Multiple MYCBP2 puncta surround the Bassoon puncta with a small region of colocalisation seen as white in the merged image. Scale bar: (A) 5 µm; (B) 0.5 µm in XY, 1 µm in Z. Images acquired on iSIM.

### iSIM super-resolution microscopy and analysis at the presynaptic bouton

Prior studies have revealed that in synuclein null mice, the volume of excitatory presynaptic boutons is decreased in excitatory neurons (Greten-Harrison et al., 2010). Furthermore, it has been firmly established that α-Syn has the capacity to modulate synaptic transmission at the presynapse (Hoffmann et al., 2021; Yoon & Munson, 2018). Therefore, our objective was to devise an experimental approach for assessing potential morphological changes in the presynaptic boutons and to explore potential modifications within the AZ. Our assay design harnessed the capabilities of the iSIM, which boasts resolution exceeding that of traditional confocal microscopes and enables antibody multiplexing. The setup of channels for our assay design required careful consideration (Figure 3). Notably, we observed a significant variation in staining patterns between MYCBP2 and ELKS immunostaining in the cultured monolayer of rat cortical neurons. MYCBP2 exhibited a pronounced increase in staining around the cell body, while ELKS typically displayed reduced staining in these areas. To ensure consistent measurements, we focused our bouton analysis exclusively on those in presumed synaptic connection with dendritic sites. This differentiation was facilitated by evaluating MYCBP2 staining before commencing analysis (Figure 3A).

**Figure 3.**
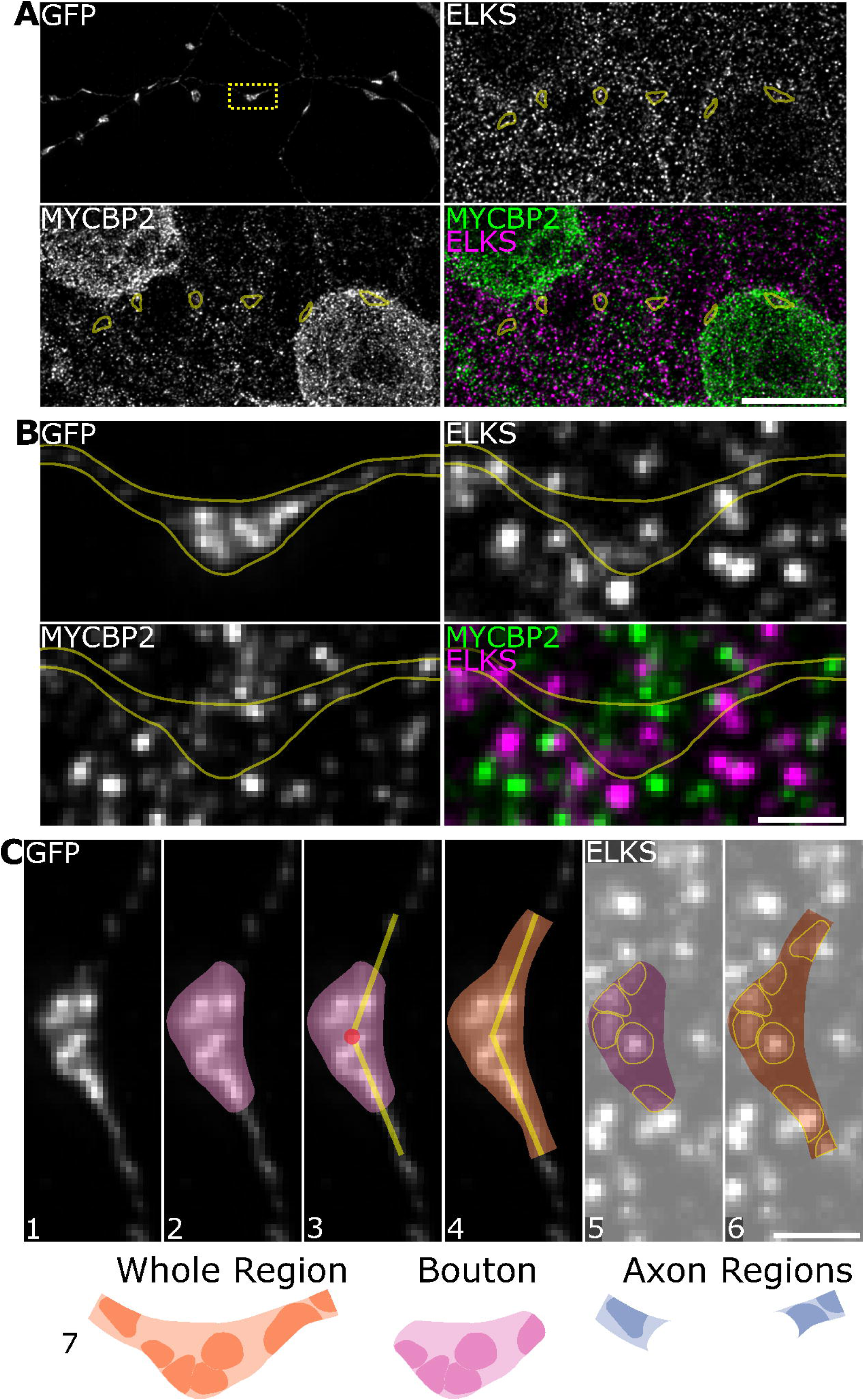
Analysis of the presynaptic bouton. **(A)** Example images of analysed regions of boutons along the axon. Rat cortical neurons were transfected with α-Syn-A30P. **(B)** An enlarged section of the inset dashed yellow box in (A) showing a single synaptic bouton. **(C)** A simplified demonstration of how the boutons and axons were analysed in ImageJ. Once the appropriate synaptic bouton is identified (1) a ROI is drawn around the bouton (2). The centre of the bouton is approximated and two lengths of 1.5 µm are drawn from the centre along the path of the axon (3). A whole ROI is then drawn (4). The bouton and the whole ROIs are overlaid on the channel of interest and the puncta are identified (5, 6). The immunostaining is then approximated in the axon regions post-acquisition by finding the difference between the whole regions and the bouton (7). Scale bar: (A) 10 µm; (B, C) 1 µm. Images acquired on iSIM.

High-resolution images revealed both MYCBP2 and ELKS staining in the synaptic bouton and adjacent axon segments (Figure 3B). Once a bouton had been selected for analysis (Figure 3C 1), a Region of Interest (ROI) was delineated around the bouton area using the GFP channel as a reference (Figure 3C 2). Our primary focus was to measure changes occurring within the presynaptic bouton. However, we also examined the adjacent axon regions to explore the possibility of proteins, such as ELKS or MYCBP2, moving between the bouton and axon regions. Given that the presynaptic boutons were often closely spaced along the axon (Figure 3A), short axon sections within 1.5 µm of each bouton’s centre were assessed (Figure 3C 3). These lengths, spanning from the bouton’s centre, marked the furthest extents of the regions to be analysed.

A ROI was created around both the bouton and the adjacent axon, collectively referred to as the ’whole’ region (Figure 3C 4). The bouton and whole regions could subsequently be utilised to measure immunoreactivity content in the respective MYCBP2 and ELKS acquisitions (Figure 3C 5&6). Following the identification of puncta within these regions, the immunoreactivity content of the axon regions was calculated post hoc. Puncta number and area in the axon regions were determined by directly comparing the whole region with the bouton region, and the intensity of axon puncta was derived from integrated intensity (Figure 3C 7). This method allowed for accurate estimation of puncta characteristics within both the bouton and the adjacent axon regions. The ImageJ script for analysis can be found online https://github.com/iainawatson/ImageJ-Analysis. The use of the iSIM provided the means to identify subtle changes and alterations in presynaptic bouton organisation, while also offering insight into protein content translocation between the bouton and axonal regions.

### Overexpression of wildtype or A30P mutant **α**-Synuclein did not alter presynaptic morphology

Prior studies have revealed that variations in α-Syn levels can influence the size of presynaptic structures (Greten-Harrison et al., 2010; Vargas et al., 2017). In our research, we aimed to explore whether the overexpression of either α-Syn-WT or the α-Syn-A30P mutant in rat cortical neurons for a duration of 3 days would lead to morphological changes in our presynaptic boutons. The transfection took place at DIV20, utilising GFP-tagged plasmids for α-Syn-WT, α-Syn-A30P, or an empty GFP vector, and the cells were allowed to express for the specified 3-day interval. Subsequently, the cells were fixed and subjected to immunostaining with α-GFP antibodies. For image processing, we deconvolved images acquired by iSIM, followed by z-stacking in ImageJ. The regions encompassing the presynaptic boutons were delineated using the GFP antibody staining. For our analysis, we selected a range of 2-4 boutons within each image, contingent upon how many boutons met our selection criteria. This approach was designed to ensure that there was no oversampling of specific cells and yielded consistent results. This process was subsequently replicated for 3-5 images per condition. Our assay for identifying the boutons within the images was highly effective. Furthermore, utilising GFP overexpressions to pinpoint the presynaptic boutons proved to be successful (Figure 4A). Upon analysing the data, we observed that the mean size of presynaptic boutons in control cells was measured at M=0.73 µm^2^, with a Standard Deviation (SD) of 0.23 µm^2^. In contrast, the mean bouton size for cells overexpressing α-Syn- WT and α-Syn-A30P was M=0.68 µm^2^ (SD=0.13 µm^2^) and M=0.74 µm^2^ (SD=0.21 µm^2^), respectively (Figure 4B). It is evident from this data that the area size of the boutons exhibited minimal variation. To substantiate these findings, a one-way ANOVA was performed, confirming that there was no significant effect of overexpressed plasmids on the bouton area at the *p* <.05 level for the three conditions (F(2,33) = 0.40, *p* = 0.6762), thus confirming our observation that the area size remained unchanged (Figure 4B).

**Figure 4.**
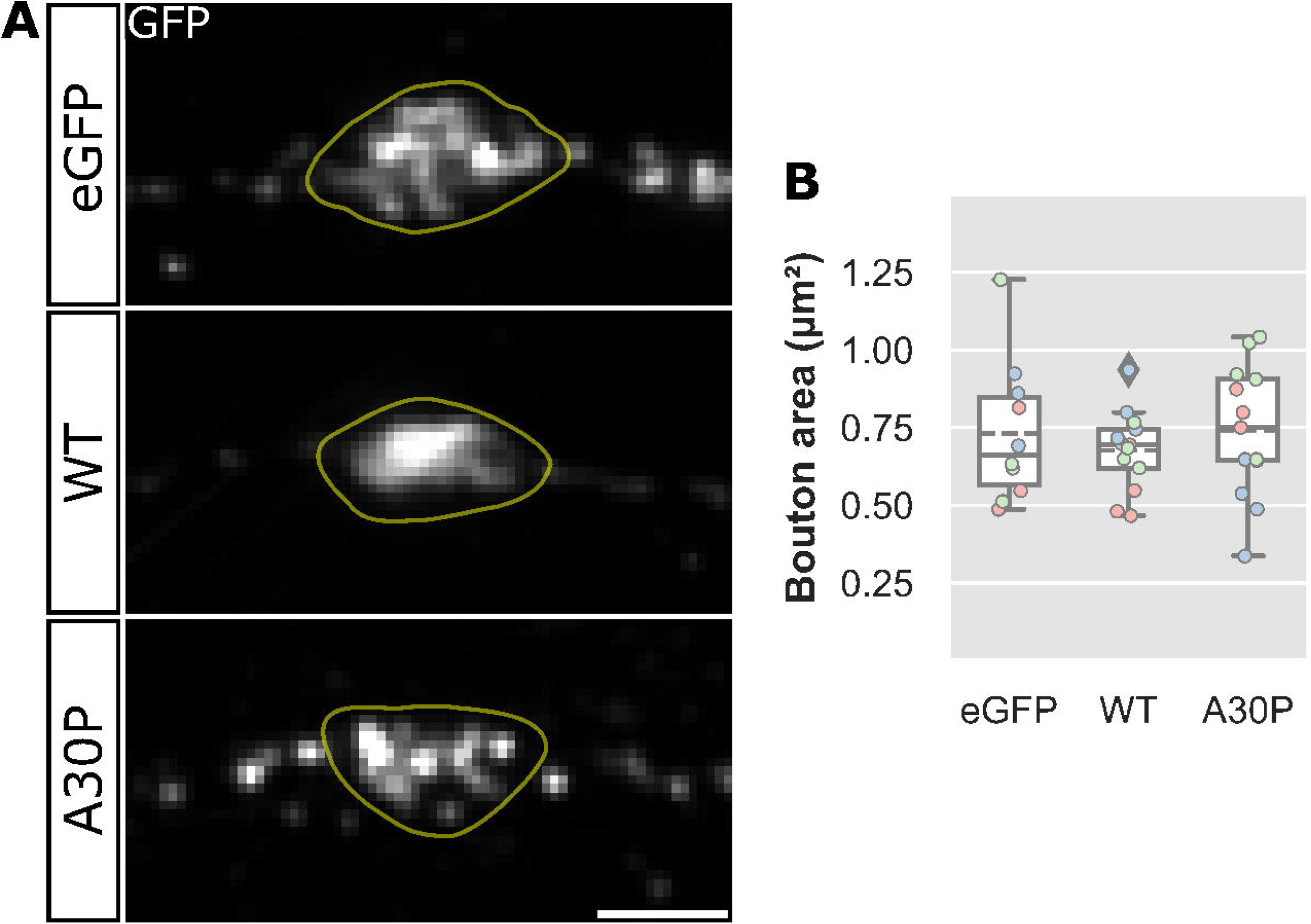
Effect of α-Synuclein on presynaptic bouton morphology. **(A)** Rat cortical neurons were transfected with eGFP, α-Synuclein-WT or α-Synuclein-A30P mutant and subsequently immunostained with GFP. The boutons were identified as indicated with the yellow lines. **(B)** No significant difference was found between any of the groups, F(2,33)=0.40, p=0.6762. Box plots: dashed lines indicate mean; colours indicate biological repeats; diamonds indicate data outside 1.5x IQR. Scale bar: 1 µm. N=2-4 boutons from 36 cells. Images acquired on iSIM.

### Overexpression of the A30P mutant **α**-Synuclein has distinctive localisation compared to wildtype **α**-Synuclein overexpression

The pathological characteristics of the α-Syn protein have been well-documented, showcasing its propensity to accumulate within the axonal processes of neurons (Braak et al., 1999). Specifically, α-Syn is known to predominantly aggregate at neuronal presynaptic terminals (Kramer & Schulz-Schaeffer, 2007). Using the iSIM we investigated for any distribution differences between cells overexpressing α-Syn- WT or α-Syn-A30P. Cultured rat cortical neurons at DIV20 were subjected to transfection with either α-Syn-GFP or α-Syn-A30P constructs, after which they were permitted to express for a duration of 3 days. Following this incubation period, the neurons underwent fixation, and immunostaining for GFP was carried out. Subsequently, the immunoreactivity of GFP was assessed, and an analysis of puncta characteristics was conducted (Figure 5A).

**Figure 5.**
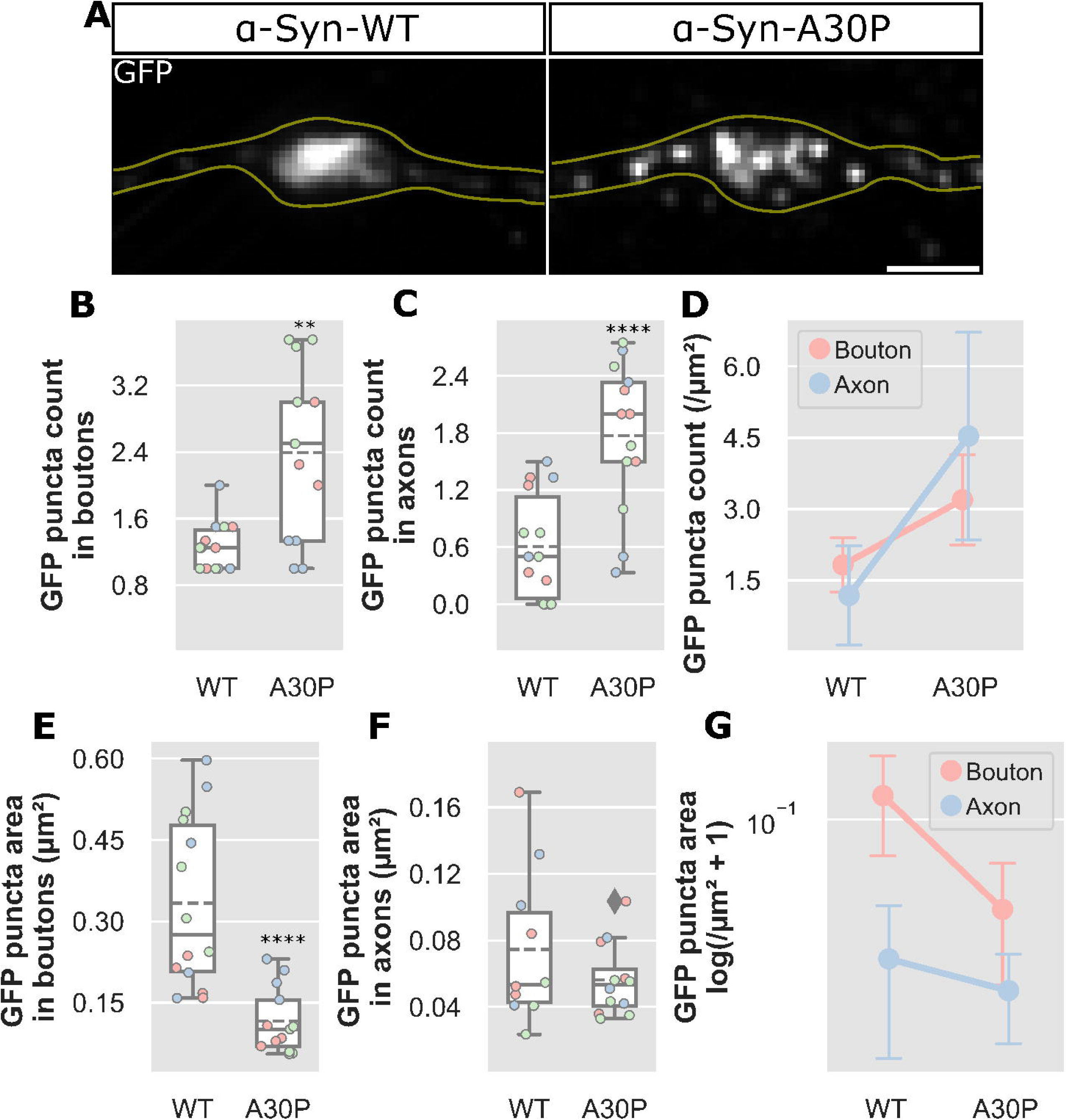
Overexpression of α-Synuclein in boutons and axons. **(A)** Rat cortical neurons were overexpressed with either eGFP tagged α-Synuclein-WT or -A30P mutant plasmids and analysed for GFP puncta count (B-D) and size (E-G). **(B, C)** In boutons, GFP tagged A30P puncta significantly increased in number compared to WT, U(31.5), p=0.0018. GFP puncta also increased significantly in adjacent axons in A30P compared to WT, t(25)=4.47, p=0.0001. **(D)** Counts were normalised to bouton area, and demonstrated a significant interaction between presynaptic regions and plasmid used, F(1.0,50.0)=7.11, p=0.0103. **(E, F)** The GFP puncta area also showed a significant reduction in A30P compared to WT within boutons, U(12.0), p=0.0001 but there was no significant change in the axon region. **(G)** Log transformed values for puncta area compared across bouton and axon regions and plasmid used demonstrated a significant interaction, F(1.0,45.0)=14.44, p=0.0004. Box plots: dashed lines indicate mean; colours indicate biological repeats; diamonds indicate data outside 1.5x IQR. Scale bar: 1 µm. Error bars: (D, G) SD. N=2-4 boutons from 25-27 cells. Images acquired on iSIM.

The quantification of GFP puncta within both the bouton and axon regions was conducted for both the GFP-tagged α-Syn-WT and -A30P overexpression constructs. Notably, both bouton and axon regions exhibited a significant increase in puncta count in the α-Syn-A30P mutant condition when contrasted with the -WT condition. The overexpressed α-Syn-A30P collected within the bouton as smaller discreet puncta, whereas the α-Syn-WT overexpression accumulated together within the bouton (Figure 5A). In the boutons, a significant difference was detected as confirmed by Mann-Whitney U test between the α-Syn-WT group (Mdn=1.25) and the -A30P condition (Mdn=2.5); *p*=0.0018, η^2^=0.313 (Figure 5B). Within the axon regions, an independent samples T-test also established a significant difference between α-Syn-A30P GFP puncta (M=1.77, SD=0.78) and α-Syn-WT GFP puncta (M=0.61, SD=0.56); t(25)=4.47, *p*=0.0001, η^2^=0.444 (Figure 5C). Since both regions were analysed separately, it was considered that the changes in puncta number might differ depending on the region. To ascertain whether this was indeed the case, a two-way ANOVA was executed to explore the impacts of region analysed and the type of α-Syn overexpression on GFP puncta count per µm^2^. The results unveiled a significant interaction between the region measured and GFP puncta overexpression; F(1.0,50.0)=7.11, *p*=0.0103, with η^2^=0.073. Consequently, it was established that the GFP puncta in the GFP-tagged α-Syn-A30P overexpression condition exhibited an increase in count in both boutons and the surrounding regions of axons when compared to α-Syn-WT overexpression. Moreover, this change in puncta count was contingent on the region measured, signifying a differential impact of α-Syn overexpression on boutons and axons concerning puncta density.

We further characterised the accumulation patterns of α-Syn-WT and α-Syn-A30P mutant by measuring puncta areas within the synaptic boutons and the surrounding axonal regions. Notably, α-Syn-A30P manifested as smaller discrete puncta within the boutons, whereas α-Syn-WT overexpression resulted in a collective accumulation within the bouton (Figure 5A). A Mann-Whitney U test validated a significant reduction in area of α-Syn-A30P puncta (Mdn=0.1) within the boutons compared to α-Syn-WT (Mdn=0.27); U(12.0), *p*=0.0001, η^2^=0.537. Conversely, the area size of α-Syn-A30P puncta (M=0.06, SD=0.02) exhibited no significant change within the surrounding axonal regions when compared to the α-Syn-WT puncta (M=0.07, SD=0.05); t(20)=1.23, *p*=0.2337, η^2^=0.017 (Figure 5F). Furthermore, a two-way ANOVA provided confirmation of a significant interaction between the region analysed and the α-Syn overexpression regarding the area size of GFP-tagged α-Syn puncta; F(1.0,45.0)=14.44, *p*=0.0004, with η^2^=0.116 (Figure 5G). In summary, α-Syn-WT overexpression in rat cortical neurons localised to presynaptic boutons and accumulated within the bouton. In contrast, α-Syn-A30P mutant overexpression also localised to presynaptic boutons, but the distribution differed from that of α-Syn-WT, forming more discrete puncta within the bouton.

### The localisation of MycBP2 is altered with overexpressed **α**-Synuclein

Recent research in *Drosophila* has revealed the interaction between α-Syn overexpression and dNmnat, which further modulates the AZ at the NMJ (Bridi et al., 2021). Moreover, dNmnat activity is subject to regulation by the E3 ligase Highwire (Russo et al., 2019). The applicability of this mechanism in mammals remained uncertain, prompting our exploration of whether α-Syn might influence MYCBP2 expression within the presynaptic boutons of rat cortical neurons. Rat cortical neurons at DIV20 were transfected with eGFP, GFP-tagged α-Syn-WT, or GFP-tagged α-Syn-A30P overexpression constructs, and the immunoreactivity of MYCBP2 was assessed. Analysis encompassed the area size, puncta number, and the intensity of MYCBP2 within the boutons and the surrounding axonal regions (Figure 6A).

**Figure 6.**
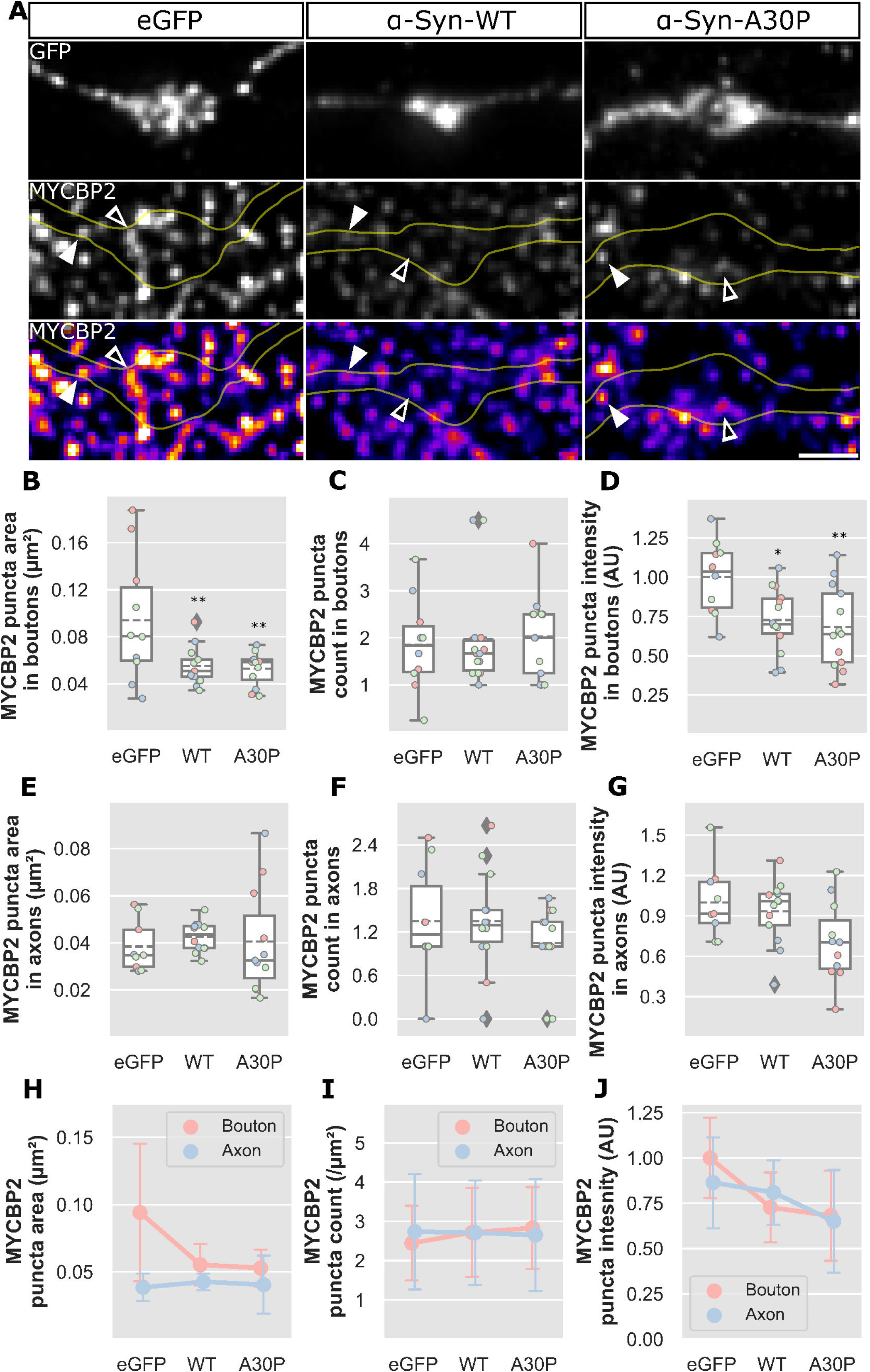
MYCBP2 localisation in response to α-Synuclein overexpression. **(A)** Rat cortical neurons were transfected with either eGFP, α-Synuclein-WT or α-Synuclein-A30P. The cells were immunostained for both GFP and MYCBP2. Hollow arrows indicate MYCBP2 lying within the bouton regions. The solid arrows show MYCBP2 puncta distributed within the axon region. The bottom pseudo-coloured panels of MYCBP2 demonstrate the signal intensity of the MYCBP2 within the above fields of view. **(B-D)** The characteristics of MYCBP2 in boutons were analysed as follows: (B) MYCBP2 puncta area significantly altered by both α-Syn-WT and -A30P mutant overexpressions compared to control F(2,32)=5.85, p=0.0068; (C) MYCBP2 count within boutons did not significantly alter; (D) Relative intensity of MYCBP2 puncta was significantly reduced in the α-Syn-WT and -A30P conditions F(2,34)=6.00, p=0.0057. **(E-G)** The characteristics of MYCBP2 in axon segments were similarly analysed for (E) area size, (F) count and (G) intensity, however, no significant differences were observed in these conditions. **(H-J)** Two-way ANOVA analysis was conducted to examine interaction effects within the following conditions: (H) There was a significant interaction between the region measured and the puncta area F(2,61)=5.09, p=0.0091; (I) Significant interaction was not identified between the region measured and the puncta count; (J) The puncta intensity did not show a significant interaction with the region measured. Box plots: dashed lines indicate mean; colours indicate biological repeats; diamonds indicate data outside 1.5x IQR. Scale bar: 1 μm. Error bars: (H-J) SD. N=2-4 boutons from 32-37 cells. Images acquired on iSIM.

Immunoreactivity for MYCBP2 was notably present within synaptic boutons (Figure 6A, hollow arrows) as well as along adjacent axons (Figure 6A, solid arrows). In the bouton, MYCBP2 puncta in the α-Syn-WT and -A30P conditions were observed to significantly reduce in area size compared to eGFP, as confirmed by one-way ANOVA at the *p*<.05 level; F(2,32)=5.85, *p*=0.0068, η^2^=0.268. Post hoc Tukey HSD comparisons demonstrated that the mean score for the eGFP condition (M=0.094, SD=0.054) significantly differed from the α-Syn-WT condition (M=0.055, SD=0.016), showing a mean difference of -0.039 µm, *p*=0.0157. Furthermore, there was a significant difference between the eGFP condition and the α-Syn-A30P condition (M=0.053, SD=0.014), with a mean difference of -0.041 µm, *p*=0.0115 (Figure 6B). However, α-Syn overexpression did not yield a significant effect on the number of MYCBP2 puncta counted within the presynaptic bouton; H(2)=0.42, *p*=0.8116 (Figure 6C).

In line with the changes in MYCBP2 area size, the intensity of MYCBP2 was also observed to significantly reduce within the presynaptic bouton at the *p*<.05 level for all three conditions (F(2,34)=6.0, *p*=0.0057, η^2^=0.261). To pinpoint the specific differences, Tukey HSD post hoc comparisons showed that the eGFP condition (M=1, SD=0.23) was significantly different from the α-Syn-WT condition (M=0.73, SD=0.20), with a mean difference of -0.27, *p*=0.0201. The α-Syn-A30P condition (M=0.68, SD=0.26) was also significantly different from the control, with a mean difference of -0.32, *p*=0.0069 (Figure 6D). It’s important to note that intensity measurements were calculated as the mean derived from the integrated intensity, thereby accommodating variations in area size.

The changes introduced by the α-Syn-WT and -A30P mutant were not mirrored in the axonal regions surrounding the presynaptic bouton. There was no change in the area size of the MYCBP2 puncta in these axonal regions (F(2,29)=0.20, *p*=0.8216) (Figure 6E). Likewise, there was no observed change in the puncta count of MYCBP2 in these axonal regions (H(2)=1.25, *p*=0.5355) (Figure 6F). However, there was a significant effect of the overexpressions on MYCBP2 puncta intensity within the axonal regions, as determined by one-way ANOVA at the *p*<.05 level (F(2,30)=3.46, *p*=0.0444) (Figure 6G). Tukey HSD post hoc comparisons did not reveal any significant differences between any of the groups, but it’s noteworthy that the mean score for the eGFP condition (M=1, SD=0.27) compared to the α-Syn- A30P condition (M=0.71, SD=0.30) demonstrated a mean difference of -0.29, *p*=0.0526.

To elucidate the interplay between α-Syn overexpression and its potential differential effects on MYCBP2 characteristics, we conducted a two-way ANOVA analysis to explore interactions. Significantly, we observed an interaction dependent on the plasmids for overexpression and the regions measured, whether bouton or axon (F(2.0,61.0)=5.09, *p*=0.0091) (Figure 6H). In contrast, there was no significant interaction between the overexpressions, and the region analysed, neither on MYCBP2 puncta count (F(2.0,68.0)=0.19, *p*=0.8272) (Figure 6I), nor on MYCBP2 puncta intensity (F(2.0,64.0)=1.36, *p*=0.2629) (Figure 6J). Collectively, the overexpression of either α-Syn-WT or -A30P mutant results in a reduction in MYCBP2 puncta size within the presynaptic bouton, with this effect exclusively observed within the bouton of α-Syn-WT overexpression. However, it is plausible that the physical constraints of the axonal regions themselves limit the size of observed puncta within these areas. Furthermore, there is a simultaneous decrease in MYCBP2 puncta intensity within the presynaptic regions, providing further evidence to suggest an overall reduction in MYCBP2 expression in response to α- Syn overexpression. This α-Syn-induced reduction in MYCBP2 is not confined to the bouton but also appears to be reduced by α-Syn-A30P predominantly within the axon. However, it’s worth noting that the results do not offer sufficient statistical significance to definitively assert the effect of α-Syn-A30P on MYCBP2 puncta intensity in the axon.

### The Overexpression of **α**-Synuclein-WT reduces the density of the presynaptic protein ELKS

The presynaptic AZ is a pivotal component of synaptic signalling transmission and serves as the site at which neurotransmitters are released (Südhof, 2012). In *Drosophila* with α-Syn overexpression, there is evidence of AZ impairment, characterised by a reduction in the levels of the protein BRP at the NMJ (Bridi et al., 2021). Therefore, our objective was to explore the impact of α-Syn overexpression on the mammalian homologue of BRP, known as ELKS. To address this question, we conducted experiments by transfecting rat cortical neurons at DIV20 with plasmids expressing eGFP, α-Syn-WT, or α-Syn-A30P. Our investigation aimed to assess the immunoreactivity and distribution of ELKS within the presynaptic boutons.

ELKS, represented by hollow arrows in Figure 7A, exhibited a discrete punctate distribution within the presynaptic bouton. Notably, there were no significant alterations in ELKS puncta area in response to the overexpression of either α-Syn- WT or α-Syn-A30P, as demonstrated by a Kruskal-Wallis test (H(2)=0.42, *p*=0.8102) (Figure 7B). Likewise, the number of ELKS puncta counted within the presynaptic bouton remained consistent across conditions, showing no significant differences according to one-way ANOVA analysis (F(2,34)=1.39, *p*=0.2620) (Figure 7C). In contrast, we observed a significant effect of the overexpression plasmids on the intensity of ELKS puncta within the presynaptic bouton, as determined by one-way ANOVA at *p*<.05; F(2,32)=5.28, *p*=0.0104, η^2^=0.248 (Figure 7D). Post hoc comparisons, conducted using the Tukey HSD method, revealed a substantial reduction in ELKS puncta intensity in both the α-Syn-WT overexpression (M=0.74, SD=0.29) with a mean difference of -0.26, *p*=0.0174, and the α-Syn-A30P overexpression (M=0.74, SD=0.13) with a mean difference of -0.26, *p*=0.0185, when compared to the eGFP control condition (M=1, SD=0.27).

**Figure 7.**
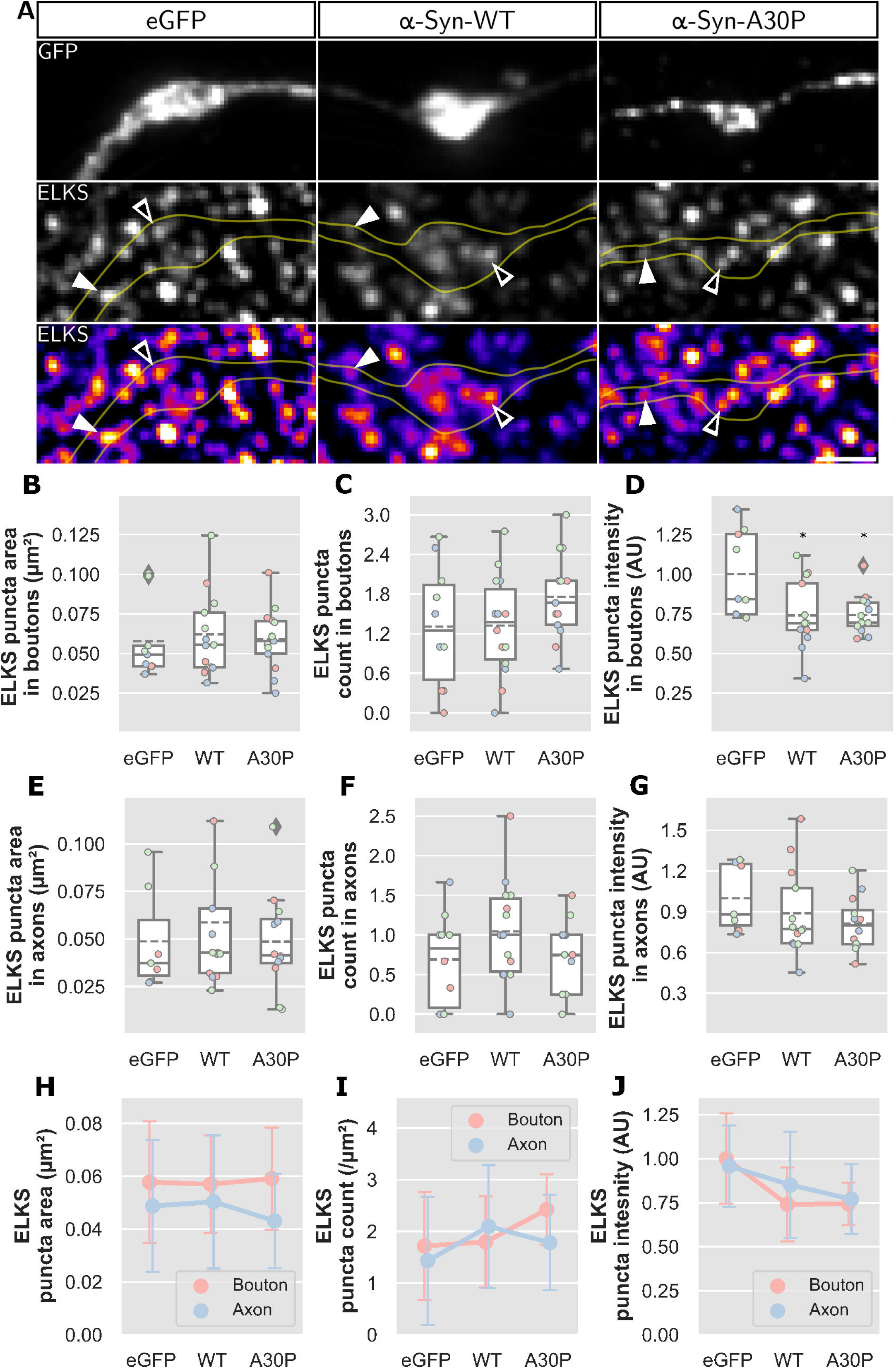
Effect of α-Synuclein overexpression of the organisation of the active zone marker ELKS. **(A)** Rat cortical neurons were transfected with either eGFP, α-Syn-WT or -A30P. The cells were immunostained for both GFP and ELKS. Hollow arrows indicate ELKS lying within the bouton regions. The solid arrows show ELKS puncta distributed within the axon region. The bottom pseudo-coloured panels of ELKS demonstrate the signal intensity of the ELKS within the above fields of view. **(B-D)** The characteristics of ELKS in boutons between overexpression conditions were analysed as follows: (B) ELKS puncta area showed no significant alterations; (C) ELKS puncta count within boutons also showed no significant change; (D) The relative intensity of ELKS in the bouton was significantly altered in the α-Syn-WT and -A30P conditions F(2,32)=5.28, p=0.0104. **(E-G)** The characteristics of ELKS in axon segments were similarly analysed for (E) area size, (F) count and (G) intensity, however, no significant differences were observed in these conditions. **(H-J)** Two-way ANOVA analysis was conducted to examine interaction effects between the area analysed and the following conditions: (H) Puncta area; (I) Puncta count; (J) Puncta intensity. No significant interaction was observed in any of these conditions. Box plots: dashed lines indicate mean; colours indicate biological repeats; diamonds indicate data outside 1.5x IQR. Scale bar: 1 μm. Error bars: (H-J) SD. N=2-4 boutons from 35-37 cells. Images acquired on iSIM.

We further analysed the characteristics of ELKS puncta within the axonal regions surrounding the bouton, focusing on the same properties identified in the boutons. However, there were no significant changes in the area size of ELKS puncta within these axonal regions (H(2)=0.7, *p*=0.7061) (Figure 7E). Additionally, there were no significant changes in the number of ELKS puncta in these regions, as determined by one-way ANOVA (F(2,34)=1.49, *p*=0.2393) (Figure 7F). In contrast to the observed effects of α-Syn overexpression within the presynaptic boutons, there was no significant impact on the intensity of ELKS puncta within the surrounding axon regions, as assessed by a Kruskal-Wallis test (H(2)=2.47, *p*=0.2901) (Figure 7G).

To explore potential interactions between the overexpression conditions and the regions analysed in terms of ELKS puncta characteristics, we employed a two-way ANOVA. Notably, the analysis revealed no significant interactive effect on ELKS puncta area (F(2.0,58.0)=0.27, *p*=0.7610) (Figure 7H), ELKS puncta count (F(2.0,68.0)=1.39, *p*=0.2567) (Figure 7I), or ELKS puncta intensity (F(2.0,61.0)=0.54, *p*=0.5883) (Figure 7J). Collectively, these findings suggest that the overexpression of α-Syn over a brief period does not have a pronounced effect on ELKS at the presynaptic terminal. The size and number of ELKS puncta remain unaltered, while a reduction in the intensity of ELKS immunoreactivity within the presynaptic bouton is indicative of a decrease in the ELKS protein content in response to α-Syn overexpression.

### Colocalisation of MYCBP2 and ELKS within presynaptic boutons and surrounding axons

We have previously examined the effects of α-Syn overexpression on the proteins ELKS and MYCBP2 at the presynapse. However, to gain a more comprehensive understanding of these effects, super-resolution microscopy may help to discern subtle alterations in protein localisation. In this study, we aimed to investigate the colocalisation of ELKS and MYCBP2 to better comprehend if they interact in response to α-Syn overexpression. We employed ELKS as a marker for the AZ of presynaptic terminals. Given that ELKS puncta number and area size remain stable following the overexpression of α-Syn-WT and -A30P, we explored whether MYCBP2 localises differentially with respect to ELKS and whether this interaction is affected by α-Syn-WT or -A30P overexpression. Cultured rat cortical neurons were allowed to mature to DIV20 before transfection with eGFP, α-Syn-WT, or α-Syn- A30P. We assessed colocalisation by identifying positive MYCBP2 intensity above background intensity within ELKS puncta ROI.

Our results revealed evident colocalisation between MYCBP2 and ELKS within the presynaptic regions of rat cortical neurons. In Figure 8A, MYCBP2 is pseudo-coloured green, and ELKS is pseudo-coloured magenta. Overlapping areas are depicted in white, as indicated by the solid white arrows within the presynaptic boutons and the hollow white arrows in the surrounding axon regions. We found that the overexpression of α-Syn-WT or -A30P did not significantly impact the colocalisation of MYCBP2 and ELKS within the boutons of our cells, as determined by one-way ANOVA (F(2,34)=0.09, *p*=0.9133) (Figure 8B). Similarly, there was no significant effect on the colocalisation within the axon regions surrounding the presynaptic bouton, also based on one-way ANOVA analysis (F(2,33)=0.76, *p*=0.4744) (Figure 8C).

**Figure 8.**
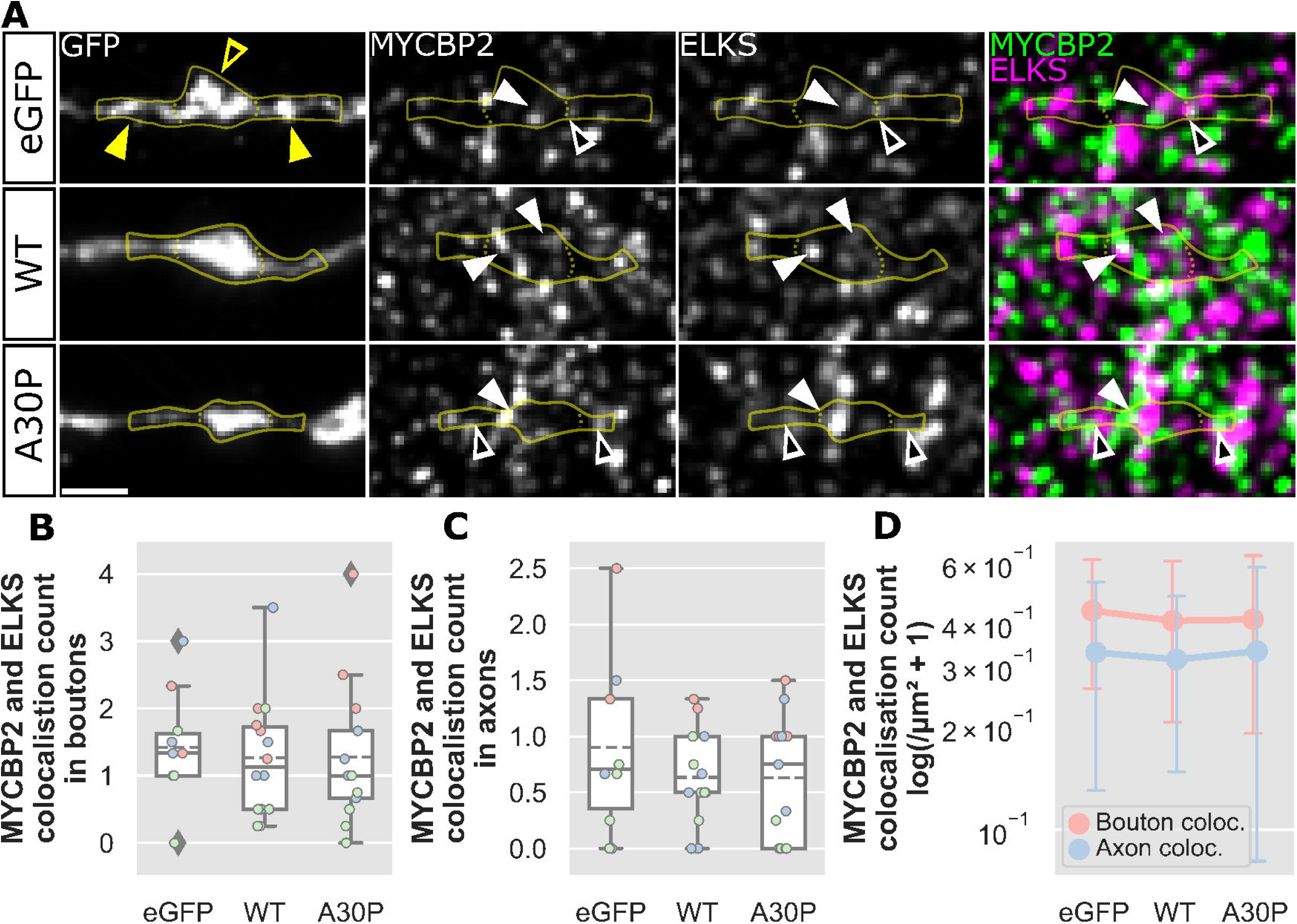
Colocalisation of MYCBP2 and ELKS and presynaptic sites. **(A)** Rat cortical neurons were transfected with either eGFP, α-Syn-WT or -A30P overexpression plasmids. The regions that were measured are identified in the GFP panel with yellow arrows. The hollow yellow arrows indicate the bouton regions analysed, and the solid yellow arrows demonstrate the axon regions analysed. The last pseudo-coloured panels show MYCBP2 immunostaining in green and ELKS immunostaining in magenta. The overlap of these colours shows as white indicating sites of colocalisation. The solid arrows indicate colocalisation within the bouton regions, whereas the hollow arrows identify colocalised puncta in the axon regions. **(B-C)** There was no significant difference to the colocalisation count of MYCBP2 and ELKS in neither the boutons (B) or the axon regions (C). **(D)** There was no significant interaction between the region analysed and the type of overexpression on the colocalisation count of MYCBP2 and ELKS. Box plots: dashed lines indicate mean; colours indicate biological repeats; diamonds indicate data outside 1.5x IQR. Scale bar: 1 µm. Error bars: (D) SD. N=2-4 boutons from 36-37 cells. Images acquired on iSIM.

Furthermore, our two-way ANOVA analysis revealed that the interaction between the region measured, and the overexpression conditions did not significantly affect the colocalisation count of MYCBP2 and ELKS (F(2.0,66.0)=0.02, *p*=0.9769) (Figure 8D). It appears that MYCBP2 interacts at the AZ region of the presynaptic bouton, as indicated by its colocalisation with ELKS. Importantly, our study demonstrates that the overexpression of α-Syn does not alter the localisation of MYCBP2 to the AZ sites of the presynaptic terminal when using ELKS as a proxy for the AZ.

## Discussion

In this study, we investigated whether elevation of α-Syn expression alters the nanoscopic organisation of the presynaptic AZ. We used iSIM super resolution microscopy to examine presynaptic bouton morphology and the distribution and organisation of AZ-associated proteins MYCBP2 and ELKS. Our findings did not reveal any pronounced changes in the morphology of presynaptic boutons resulting from the overexpression of α-Syn. However, a marked distinction emerged in the distribution of α-Syn-WT and α-Syn-A30P overexpressions, with the former exhibiting a notably more condensed aggregation within the boutons. Examination of MYCBP2 within the presynapse, revealed that overexpression of either α-Syn-WT or α-Syn-A30P led to a reduction in puncta size and MYCBP2 intensity. Furthermore, we observed a decrease in ELKS immunoreactivity without any concurrent alterations in puncta size. Collectively, these findings suggest that the overexpression of α-Syn can manifest its effects through a mechanism implicating MYCBP2 and ELKS. These presynaptic changes unveil a novel avenue through which α-Syn may exert its influence within the presynapse, shedding light on potential mechanisms within synucleinopathies like PD.

### The expression profile and morphological impact of **α**-Synuclein

The accumulation of α-Syn within presynaptic terminals is a recognised feature of synucleinopathies, including Dementia with Lewy Bodies (DLB) (Kramer & Schulz-Schaeffer, 2007). In post-mortem tissue, α-Syn aggregates are frequently observed in presynaptic regions, distinct from Lewy body inclusions, and are accompanied by reduced pre- and postsynaptic markers and dendritic spine loss. These findings indicate that α-Syn accumulation can directly disrupt presynaptic structure and connectivity (Kramer & Schulz-Schaeffer, 2007). However, the underlying mechanisms and temporal sequence of these changes remain unclear. Animal studies have shown that the physiological functions of α-, β-, and γ-synucleins overlap, complicating interpretation of single-gene manipulations. While triple-knockout (αβγ-Syn-/-) mice exhibit reduced bouton size (Vargas et al., 2017), single or double knockouts do not, likely due to compensatory upregulation of the remaining synucleins (Chandra et al., 2004). This redundancy suggests that presynaptic morphology is resilient to partial disruption of synuclein expression. In this context, our finding that α-Syn overexpression did not alter bouton morphology aligns with the idea that structural changes at the presynapse arise gradually or require more extensive perturbation, such as chronic accumulation or loss of all synuclein isoforms.

Methodologically, measuring bouton size even with super resolution imaging, presents inherent challenges. We defined bouton boundaries using GFP fluorescence, saturating the signal to define edges consistently across conditions. The aggregated nature of α-Syn-WT required lower acquisition parameters to avoid image saturation. However, as shown in Figure 3, both eGFP and α-Syn-A30P overexpressions exhibit a punctate nature, which makes the precise shape and morphology somewhat estimated. This punctate distribution contrasts with the typical expression of diffuse GFP within the soma and dendrites, as demonstrated in Figure S1B. This punctate pattern likely reflects the constrained organisation of axons and boutons, where diffusion of cytosolic fluorophores is limited. While alternative markers, such as phalloidin-labelled F-actin, could in principle define presynaptic bouton outlines, these approaches typically lack cell-type specificity in mixed cultures and may obscure synaptic boundaries. Even with the use of super-resolution techniques, the identification of presynaptic actin organisation remains challenging, primarily due to occlusion by postsynaptic signals (Papandréou & Leterrier, 2018). Electron microscopy is a potential alternative method that could provide precise measurements of presynaptic morphology (Korobova & Svitkina, 2010; Vargas et al., 2017). However, it would introduce challenges related to identifying neurons solely expressing α-Syn and necessitate alternative methods such as the use of α-Syn overexpression animal models, which are beyond the scope of our experimental design. We therefore interpret bouton size measurements as relative, rather than absolute, but robust across matched conditions.

### The distribution pattern of overexpressed **α**-Synuclein at presynaptic boutons

The distinct distribution patterns of α-Syn WT and A30P observed may reflect their known differences in membrane association and aggregation behaviour. The A30P mutation, a rare autosomal dominant variant linked to familial PD (Krüger et al., 1998), reduces α-Syn’s affinity for lipid membranes (Cole et al., 2002; Fortin et al., 2004; Kubo et al., 2005). Consequently, α-Syn-A30P displays a more diffuse distribution within boutons, whereas α-Syn-WT tends to form denser, more compact accumulations. Previous work has shown that α-Syn-WT overexpression impairs synaptic vesicle exocytosis (Larsen et al., 2006; Lundblad et al., 2012; Nemani et al., 2010). These effects have been attributed to alterations in vesicle clustering or late-stage exocytosis (Larsen et al., 2006; Nemani et al., 2010). On the other hand, the α- Syn-A30P variant has been demonstrated to exert differing effects on depolarisation-evoked dopamine release and presynaptic vesicle density compared to α-Syn-WT (Gaugler et al., 2012). Despite these distinct effects on presynaptic function, both α-Syn variants may ultimately converge on similar downstream disruptions when expressed at high levels (Larsen et al., 2006). Consistent with this idea, we observe that both α-Syn-WT and α-Syn-A30P reduce MYCBP2 expression levels . Given that A30P is more cytosolic, its wider diffusibility could extend its effects beyond boutons, potentially explaining the broader reduction in MYCBP2 observed along axons. The denser α-Syn-WT aggregates, conversely, may act locally within boutons, where α- Syn is known to interact with synaptic vesicle membranes and other presynaptic components.

The presence of dense α-Syn-WT within the presynaptic terminal introduces an intriguing paradox. Studies have demonstrated that fibrillar α-Syn acts as a seed and actively recruits endogenous α-Syn into aggregating structures (Bousset et al., 2013). This phenomenon has also been observed in primary neuronal mouse cultures *in vitro* (Courte et al., 2020). Bousset *et al*. also found that, while α-Syn- A30P can form fibrils similar to α-Syn-WT, it cannot form ribbons, demonstrating divergent characteristics (Bousset et al., 2013). In our experimental design, it’s possible that overexpressed α-Syn may aggregate and recruit endogenous α-Syn, potentially leading to a reduction in the endogenous functionality of α-Syn. The resulting changes to MYCBP2 and ELKS may be a consequence of decreased endogenous α-Syn functionality. Loss of Synuclein cannot fully explain synucleinopathies however. For example, studies involving synuclein triple knockout mice have demonstrated no overt phenotype, no change in midbrain dopaminergic neurons, and no neurodegeneration or Parkinsonism-like phenotype (Anwar et al., 2011). There is still ongoing debate within the field about whether the pathogenic nature of α-Syn relates to increased or decreased levels of α-Syn. Future studies building upon our work should consider incorporating knockdown experiments of α-Syn. This approach would help ascertain whether reduced endogenous α-Syn would elicit similar effects on MYCBP2 and ELKS as observed with overexpressed α-Syn. Additionally, it would be invaluable to immunostain for endogenous α-Syn, and determine whether there are localisation changes associated with the endogenous protein between controls and the overexpression of α-Syn. Such investigations could shed further light on whether accumulated α-Syn drives a toxic gain-of-function or a loss-of-function at the presynapse, or perhaps even both.

### Potential consequences of **α**-Synuclein reducing MYCBP2 expression

MYCBP2, a ubiquitin E3 protein ligase, has been shown to play a pivotal role in regulating various cellular processes, including cytoskeletal dynamics, neural development, and axonal degeneration (Grill et al., 2016; Guo et al., 1998; Virdee, 2022). In general, ubiquitin E3 ligases primarily function by recruiting E2 ubiquitin-conjugating enzymes, which facilitate the transfer of ubiquitin to specific protein substrates, ultimately marking them for degradation. In *Drosophila*, Highwire is implicated in regulating the size and activity of presynaptic terminals at the NMJ (Russo et al., 2019). However, our experiments involving α-Syn overexpression in neuronal cultures did not result in changes to the AZ morphology upon the reduction of MYCBP2 levels with respect to ELKS. It is essential to note that Russo and colleagues also demonstrated that reduced levels of MYCBP2 led to an excess of dNmnat, eventually resulting in dysregulation of BRP at the AZ (Russo et al., 2019). These findings may suggest potential variations in the underlying mechanisms governing synaptic structure and regulation between different synaptic types and organisms. It seems unlikely that ELKS expression levels are being directly regulated by MYCBP2, at least in terms of ubiquitination mechanics. Therefore, MYCBP2 could be negatively regulating the mammalian homologue of dNmnat, NMNAT2, and potentially influence the dynamics of the AZ.

Highwire’s role in regulating presynaptic physiology has also been associated with a reduction in synaptic strength at the NMJ (C. Wu et al., 2005). Additionally, investigations of the *C. elegans* orthologue of MYCBP2, known as RPM-1, have revealed its necessity in the internalisation of AMPA receptors through the inactivation of p38 MAPK signalling pathways (Nakata et al., 2005). Conversely, studies in mice focusing on MYCBP2 within thermoreceptor neurons reported an increased response to nociceptor pain behaviour, indicating enhanced neurotransmission (Ehnert et al., 2004). Furthermore, reduced MYCBP2 levels in mice have also been linked to the blocking of the internalisation of transient receptor potential vanilloid receptor 1 in sensory neurons, also through p38 MAPK pathways (Holland et al., 2011). In future experiments, it would be vital to ascertain whether the reduced levels of MYCBP2 at the presynapse lead to constitutive activation of p38 MAPK.

### **α**-Synuclein induced changes at the Active Zone

Previous research by Bridi *et al*. demonstrated in *Drosophila* that overexpression of α-Syn resulted in a significant reduction in both BRP puncta size and puncta count, indicating AZ alterations (Bridi et al., 2021). However, we did not observe changes in the size or count of ELKS puncta. Instead, we detected a significant decrease in the overall immunoreactivity, indicating reduced protein density rather than structural disassembly. These differences likely reflect evolutionary divergence in AZ organisation. In *Drosophila*, BRP is essential for T-bar formation and AZ maturation (Fouquet et al., 2009; Kittel et al., 2006; Liu et al., 2011). In contrast, ELKS is only a partial homologue of BRP, sharing similarity primarily in the N-terminal section (Wagh et al., 2006). The C-terminal portion of BRP more closely resembles cytoskeletal proteins that confers its structural role. In vertebrates, this function is distributed among additional scaffolding proteins such as Bassoon and Piccolo (Dani et al., 2010; Wagh et al., 2006; Wong et al., 2018). Therefore, the differences we observed compared to the findings by Bridi *et al*. are likely due to phylogenetic disparities between model organisms; the structural and functional distinctions in the AZs of *Drosophila* and vertebrate neurons could account for these variations.

While overexpression of α-Syn variants reduced ELKS loss but did not alter bouton or AZ morphology, it may still affect presynaptic physiology. ELKS is known to regulate neurotransmitter release by controlling the size of the readily releasable vesicle pool (Held et al., 2016). Given that α-Syn modulates vesicle dynamics and that MYCBP2 can influence protein turnover and translation, reduced ELKS expression could represent a downstream effect of α-Syn-induced disruption in presynaptic signalling or local proteostasis. Therefore, future studies should focus on assessing the functional consequences of reduced ELKS, rather than further structural endpoints. Electrophysiological assays, including the use of a hypertonic sucrose solution to release the entire readily releasable pool of vesicles (Rosenmund & Stevens, 1996) could be used to determine whether α-Syn overexpression-induced reduction of ELKS expression can impact vesicle priming or release probability.

### Limitations

While this study provides valuable insights into the mechanisms underlying presynaptic mechanisms, overexpression of constructs may not recapitulate native or pathophysiological protein levels, risk non-physiological interactions and signalling artefacts. Use of fluorescent tags also raises the possibility of tag-related aggregation (e.g., for GFP β-barrel proteins), which could confound intensity-based readouts (Antonets et al., 2020). However, in such circumstances a breakdown in protein structure is necessary and is therefore exhibits much reduced fluorescence, partly mitigating this concern. In addition, conclusions are based primarily on iSIM imaging. Future studies would benefit from complementary modalities and functional assays (e.g., electrophysiology/biochemistry). Such methods would help validate the observed MYCBP2/ELKS changes and clarify how elevated α-Syn perturbs AZ composition while overall bouton morphology appears preserved.

### Summary

In this study, we investigated the impact of α-Syn on presynaptic terminals, focusing on morphological changes and potential mechanisms associated with α-Syn at the presynapse. While our findings did not reveal significant alterations in presynaptic bouton morphology due to α-Syn overexpression, we observed distinct differences in the distribution patterns of α-Syn-WT and α-Syn-A30P, with the latter showing more diffuse expression within boutons. Both α-Syn-WT and α-Syn-A30P overexpression resulted in a reduction in MYCBP2 puncta count and intensity within the presynapse. Additionally, we found a decrease in ELKS immunoreactivity without concurrent changes in puncta size. These results suggest that overexpressed α-Syn may exert its effects through mechanisms involving MYCBP2 and ELKS, shedding light on potential mechanisms within synucleinopathies like PD. Notably, while no overt structural changes were observed in the AZ, the functional consequences of reduced ELKS expression and potential alterations in synaptic dynamics merit further investigation to better understand the role of α-Syn at the presynapse. The study extends our understanding of the intricate interactions between α-Syn overexpression and presynaptic components and highlights areas for future research to explore functional changes and their implications in the context of neurodegenerative diseases.

## Abbreviations

AD: Alzheimer’s Disease
α-Syn: α-Synuclein
AZ: Active Zone
BRP: Bruchpilot
DIV: Days In Vitro
DLB: Dementia with Lewy Bodies
eGFP: enhanced GFP
HD: Huntington’s Disease
IQR: Interquartile Range
iSIM: instant Structured Illumination Microscope
NMJ: Neuromuscular Junction
PD: Parkinson’s Disease
RIM-BPs: Rab3-interacting molecule-binding proteins
ROI: Region of Interest
SD: Standard Deviation

## Author Contributions

Conceptualization, I.W., J.B., F.H. and D.P.S.; methodology, I.W., J.B. and D.P.S.; investigation, I.W.; formal analysis, I.W.; resource provision, D.H.; writing: original draft preparation, I.W.; writing: review and editing, I.W., and D.P.S.; visualisation, I.W.; resources, D.H., D.P.S.; supervision, D.P.S.; project administration, D.P.S; funding acquisition, D.P.S.

## Acknowledgements

Authors would like to thank George Chennell of the Wohl Cellular Imaging Centre (King’s College London) for technical support.

