## Supplemental Figures for "Overexpression of α-Synuclein Alters The Nanoscopic Organisation of Presynaptic Proteins"

### Nanoscopic Organisation of Presynaptic α-Synuclein in Parkinson's Disease

#### SUPPLEMENTAL FIGURES

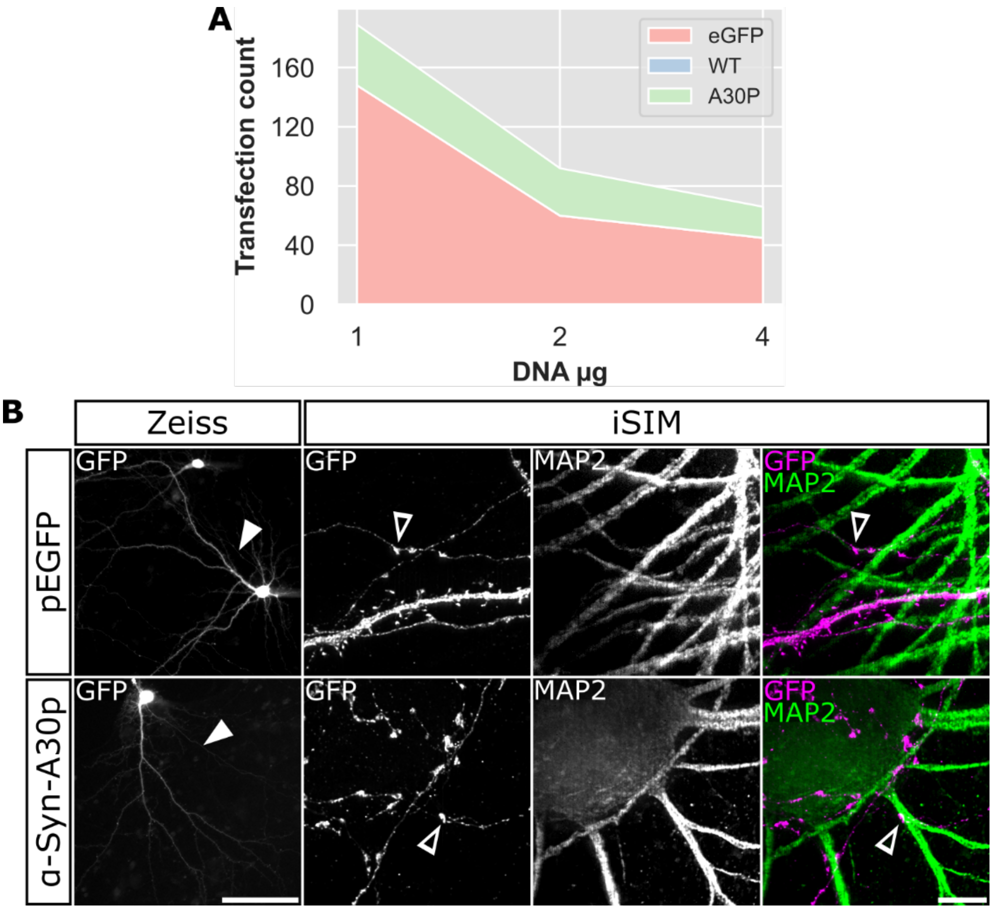


**Figure S1 α-Syn transfections in cortical neurons and synaptic boutons. (A)** α-Syn plasmids were tested for their transfection efficiency and efficacy and compared to eGFP. α-Syn plasmids include: α-Syn-Wildtype (WT); α-Syn-A30P (A30P). Transfection tests were made using a range of DNA quantity from 1 to 4 µg. Total number of transfected cells at 1, 2 and 4 µg were 189, 92 and 96 respectively. WT cells did not transfect during initial tests. **(B)** Cortical neurons were transfected with eGFP or A30P. Cells were imaged with either the Zeiss epifluorescence or iSIM. White arrowheads in the left most panel indicate the axons. Remaining panels demonstrate synaptic boutons shown with hollow arrowheads. Using MAP2 to identify dendrites, synaptic boutons are visibly associating with dendrites and regions around the cell bodies indicated by hollow arrowheads in the right panels. Zeiss scale bar 100 µm. iSIM scale bar 5 µm.

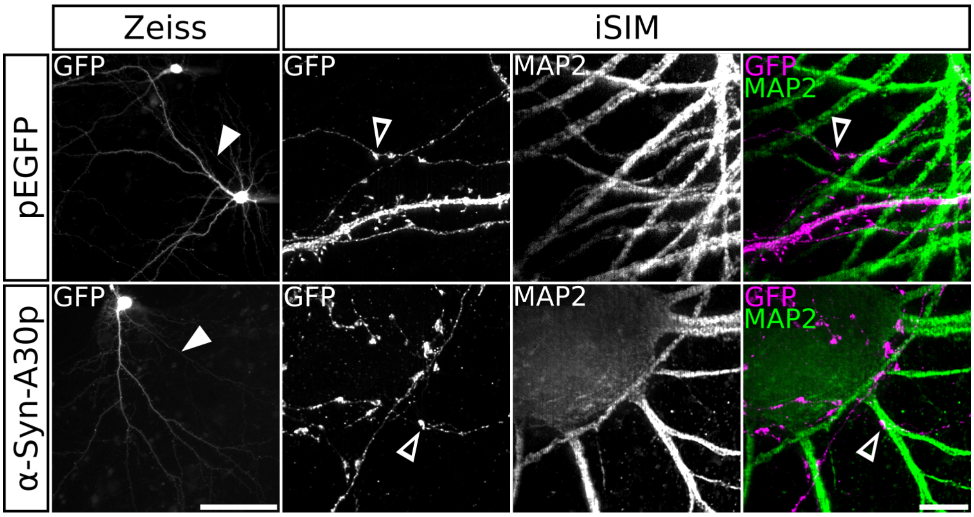
Zeiss
iSIM
pEGFP
α-Syn-A30p
GFP
GFP
MAP2
GFP
MAP2
GFP
GFP
MAP2
GFP
MAP2

**Figure 2 Transfected neurons and presynaptic boutons.** Rat cortical neurons were transfected with eGFP or α-Syn-A30P. Cells were imaged with either the Zeiss epifluorescence or iSIM. White arrowheads in the left most panel indicate the axons. Remaining panels demonstrate the presence of synaptic boutons shown with hollow arrowheads. Using MAP2 to identify dendrites, synaptic boutons are visibly associating with dendrites and regions around the cell bodies indicated by hollow arrowheads in the right panels. Zeiss scale bar 100 µm. iSIM scale bar 5 µm.

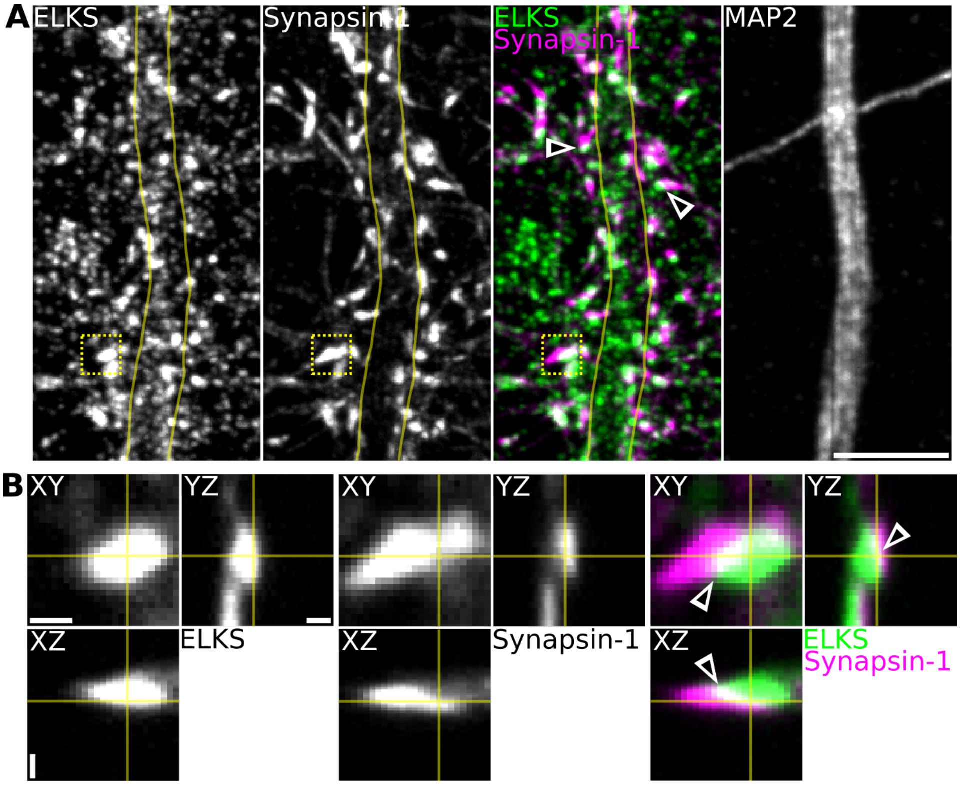
A
ELKS
Synapsin-1
ELKS
Synapsin-1
MAP2
B
XY
YZ
XZ
ELKS
XY
YZ
XZ
Synapsin-1
XY
YZ
XZ
ELKS
Synapsin-1

**Figure 3 ELKS protein distribution. (A)** Z-stacked images show sections of primary rat cortical neurons that were stained with the AZ marker ELKS. Cells were also immunostained with Synapsin-1 to confirm the presynapse and MAP2 to localise staining along sections of dendrite. In the colour merged images shades of white indicate colocalisation between ELKS and Synapsin-1, highlighted by arrows. **(B)** Orthogonal views are enlarged regions taken from the dashed yellow boxes indicated in (A) to show depth of the acquired stacks. The region of colocalisation is also given by arrows. Scale bar: (A) 5 µm; (B) 0.5 µm in XY, 1 µm in Z. Images acquired on iSIM.

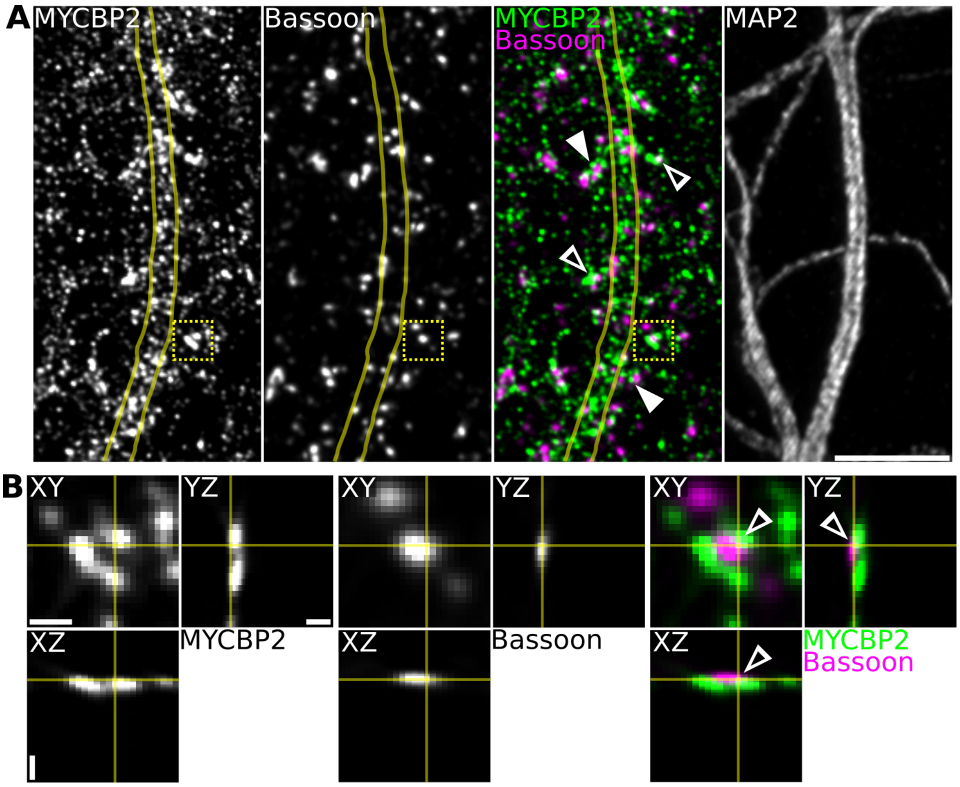
A
MYCBP2
Bassoon
MYCBP2
Bassoon
MAP2
B
XY
YZ
XZ
MYCBP2
XY
YZ
XZ
Bassoon
XY
YZ
XZ
MYCBP2
Bassoon

**Figure 4 Distribution of MYCBP2 with the presynaptic marker Bassoon. (A)** Z-stack images of rat cortical neurons immunostained for MYCBP2. MYCBP2 was coimmunostained with the presynaptic marker Bassoon to help identify presynaptic localisation. Cells were also coimmunostained with MAP2 to highlight areas of dendrite. In the merged image MYCBP2 can frequently be observed to associate closely with Bassoon as indicated by the solid arrows. Additionally, several areas are observed to colocalise as white in the merged image indicated by the hollow arrows. **(B)** Orthogonal views of the inset dashed yellow boxes in (A) to show the depth of the acquired images. Multiple MYCBP2 puncta surround the Bassoon puncta with a small region of colocalisation seen as white in the merged image. Scale bar: (A) 5 µm; (B) 0.5 µm in XY, 1 µm in Z. Images acquired on iSIM.

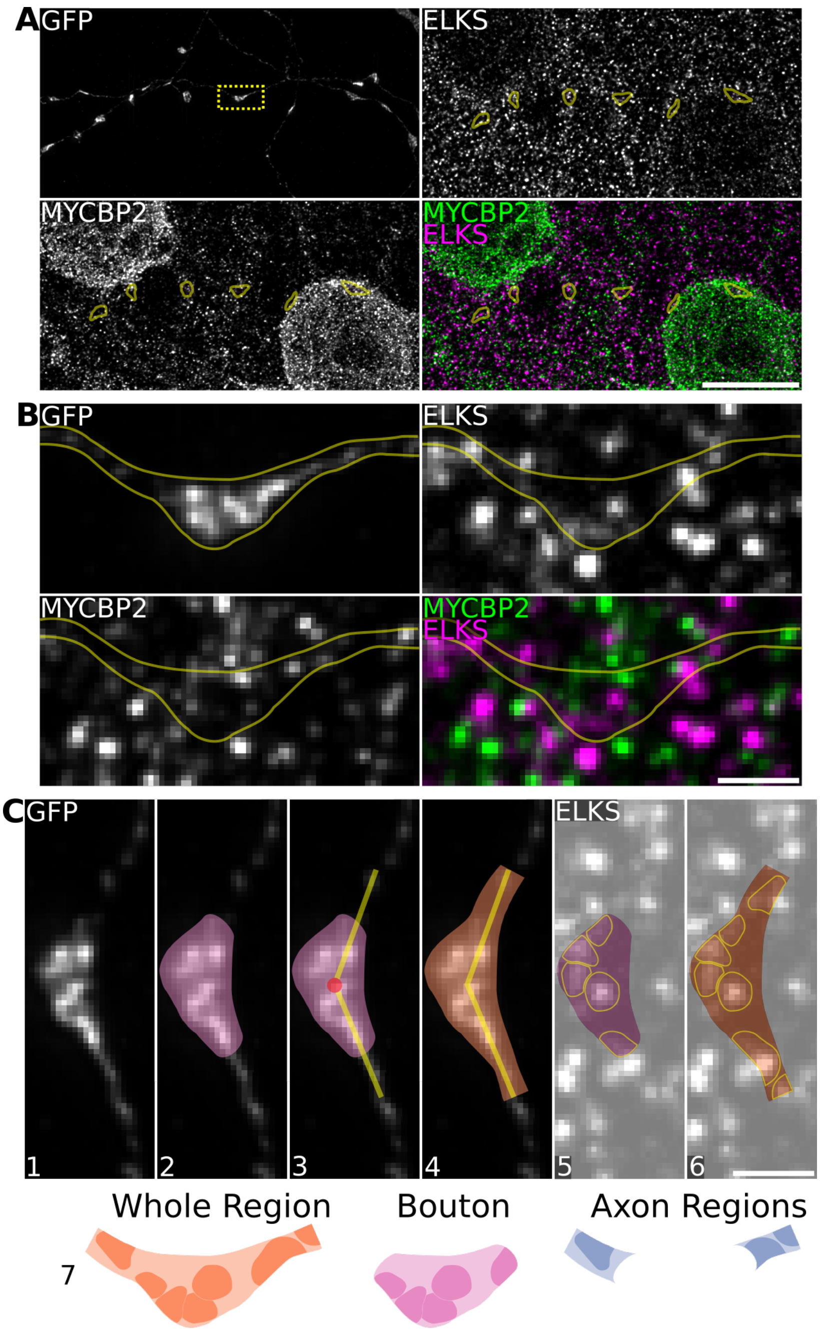
A
GFP
ELKS
MYCBP2
MYCBP2
ELKS
B
GFP
ELKS
MYCBP2
MYCBP2
ELKS
C
GFP
ELKS
1
2
3
4
5
6
Whole Region
Bouton
Axon Regions
7

**Figure 5 Analysis of the presynaptic bouton. (A)** Example images of analysed regions of boutons along the axon. Rat cortical neurons were transfected with α-Syn-A30P. **(B)** An enlarged section of the inset dashed yellow box in (A) showing a single synaptic bouton. **(C)** A simplified demonstration of how the boutons and axons were analysed in ImageJ. Once the appropriate synaptic bouton is identified (1) a ROI is drawn around the bouton (2). The centre of the bouton is approximated and two lengths of 1.5 µm are drawn from the centre along the path of the axon (3). A whole ROI is then drawn (4). The bouton and the whole ROIs are overlaid on the channel of interest and the puncta are identified (5, 6). The immunostaining is then approximated in the axon regions post-acquisition by finding the difference between the whole regions and the bouton (7). Scale bar: (A) 10 µm; (B, C) 1 µm. Images acquired on iSIM.

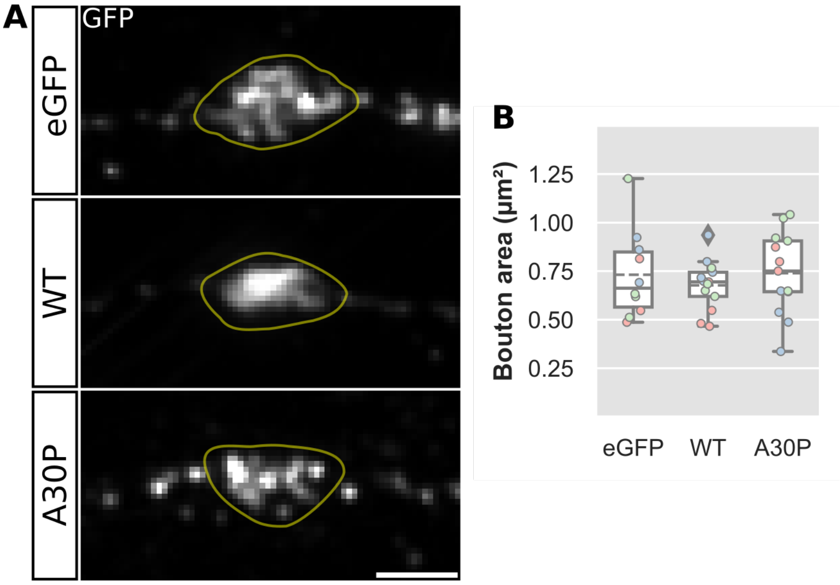
A
GFP
eGFP
WT
A30P
B
Bouton area (μm²)
1.25
1.00
0.75
0.50
0.25
eGFP
WT
A30P

**Figure 6 Effect of α-Synuclein on presynaptic bouton morphology. (A)** Rat cortical neurons were transfected with eGFP, α-Synuclein-WT or α-Synuclein-A30P mutant and subsequently immunostained with GFP. The boutons were identified as indicated with the yellow lines. **(B)** No significant difference was found between any of the groups, $F(2,33)=0.40$, $p=0.6762$. Box plots: dashed lines indicate mean; colours indicate biological repeats; diamonds indicate data outside 1.5x IQR. Scale bar: 1 µm. N=2-4 boutons from 36 cells. Images acquired on iSIM.

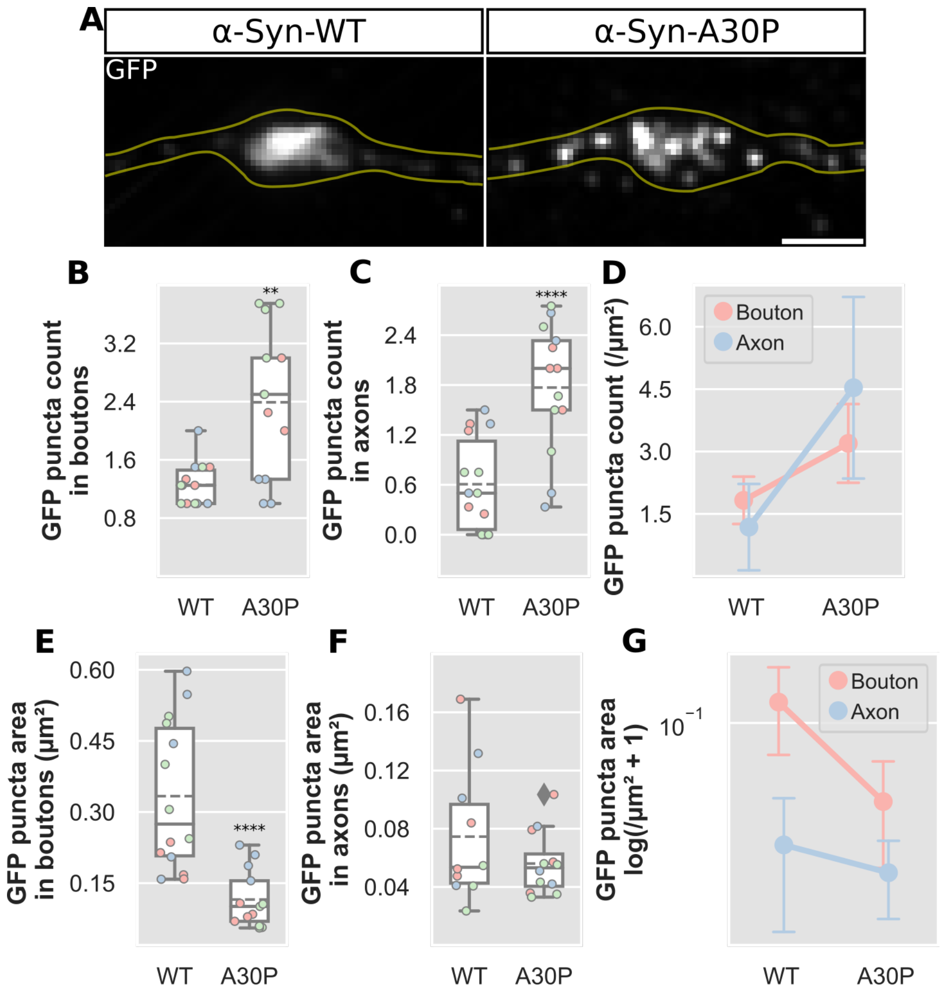

A
α-Syn-WT
α-Syn-A30P
GFP
B
GFP puncta count in boutons
3.2
2.4
1.6
0.8
**
WT
A30P
C
GFP puncta count in axons
2.4
1.8
1.2
0.6
0.0
****
WT
A30P
D
GFP puncta count (/µm²)
6.0
4.5
3.0
1.5
Bouton
Axon
WT
A30P
E
GFP puncta area in boutons (µm²)
0.60
0.45
0.30
0.15
****
WT
A30P
F
GFP puncta area in axons (µm²)
0.16
0.12
0.08
0.04
WT
A30P
G
GFP puncta area log(/µm² + 1)
10⁻¹
Bouton
Axon
WT
A30P

**Figure 7 Overexpression of α-Synuclein in boutons and axons. (A)** Rat cortical neurons were overexpressed with either eGFP tagged α-Synuclein-WT or -A30P mutant plasmids and analysed for GFP puncta count (B-D) and size (E-G). **(B, C)** In boutons, GFP tagged A30P puncta significantly increased in number compared to WT, U(31.5), p=0.0018. GFP puncta also increased significantly in adjacent axons in A30P compared to WT, t(25)=4.47, p=0.0001. **(D)** Counts were normalised to bouton area, and demonstrated a significant interaction between presynaptic regions and plasmid used, F(1.0,50.0)=7.11, p=0.0103. **(E, F)** The GFP puncta area also showed a significant reduction in A30P compared to WT within boutons, U(12.0), p=0.0001 but there was no significant change in the axon region. **(G)** Log transformed values for puncta area compared across bouton and axon regions and plasmid used demonstrated a significant interaction, F(1.0,45.0)=14.44, p=0.0004. Box plots: dashed lines indicate mean; colours indicate biological repeats; diamonds indicate data outside 1.5x IQR. Scale bar: 1 µm. Error bars: (D, G) SD. N=2-4 boutons from 25-27 cells. Images acquired on iSIM.

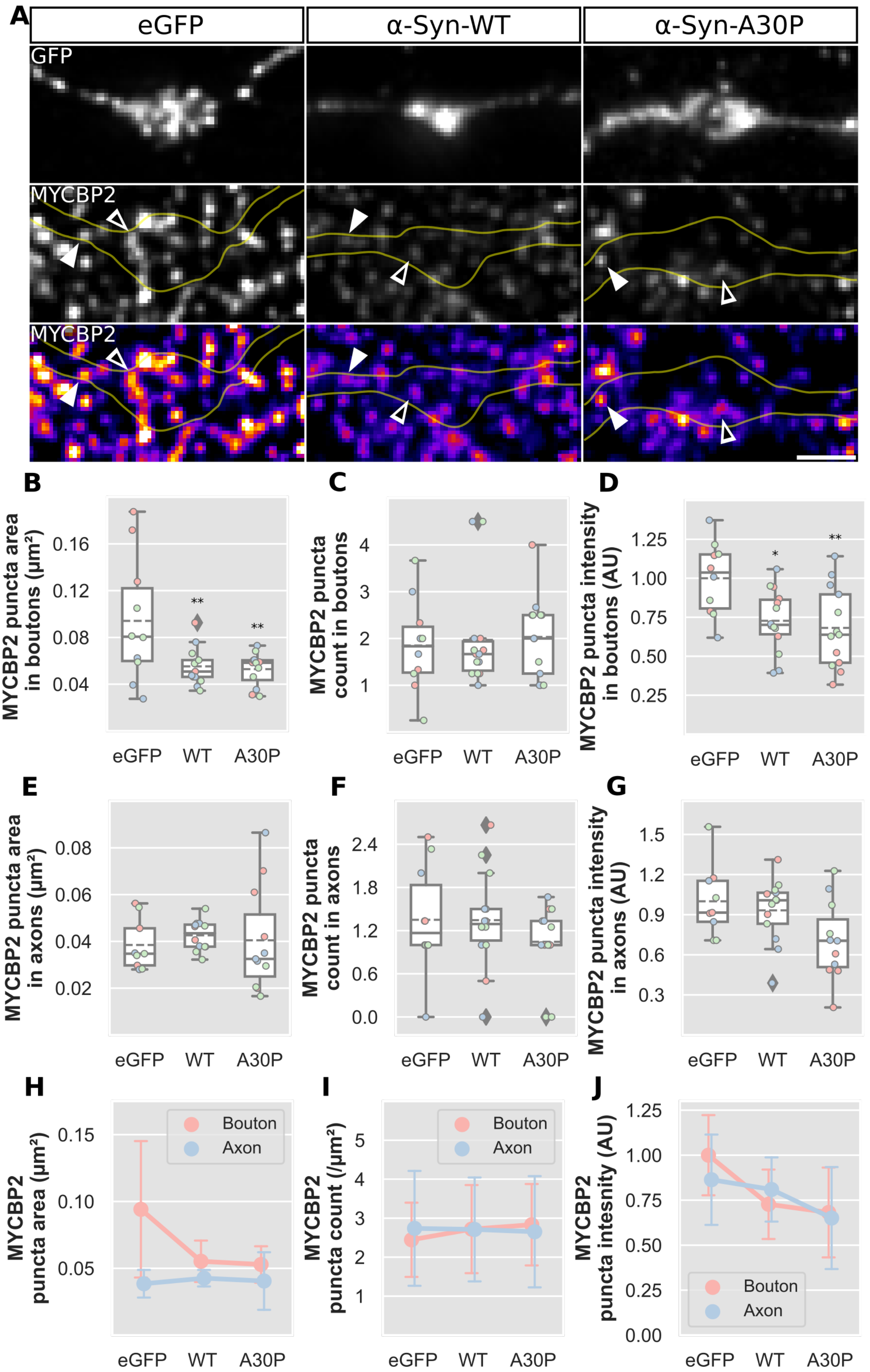

A
eGFP
α-Syn-WT
α-Syn-A30P
GFP
MYCBP2
MYCBP2
B
MYCBP2 puncta area in boutons (µm²)
0.16
0.12
0.08
0.04
**
**
eGFP
WT
A30P
C
MYCBP2 puncta count in boutons
4
3
2
1
eGFP
WT
A30P
D
MYCBP2 puncta intensity in boutons (AU)
1.25
1.00
0.75
0.50
0.25
*
**
eGFP
WT
A30P
E
MYCBP2 puncta area in axons (µm²)
0.08
0.06
0.04
0.02
eGFP
WT
A30P
F
MYCBP2 puncta count in axons
2.4
1.8
1.2
0.6
0.0
eGFP
WT
A30P
G
MYCBP2 puncta intensity in axons (AU)
1.5
1.2
0.9
0.6
0.3
eGFP
WT
A30P
H
MYCBP2 puncta area (µm²)
0.15
0.10
0.05
Bouton
Axon
eGFP
WT
A30P
I
MYCBP2 puncta count (/µm²)
5
4
3
2
1
Bouton
Axon
eGFP
WT
A30P
J
MYCBP2 puncta intesnity (AU)
1.25
1.00
0.75
0.50
0.25
0.00
Bouton
Axon
eGFP
WT
A30P

**Figure 8 MYCBP2 localisation in response to α-Synuclein overexpression. (A)** Rat cortical neurons were transfected with either eGFP, α-Synuclein-WT or α-Synuclein-A30P. The cells were immunostained for both GFP and MYCBP2. Hollow arrows indicate MYCBP2 lying within the bouton regions. The solid arrows show MYCBP2 puncta distributed within the axon region. The bottom pseudo-coloured panels of MYCBP2 demonstrate the signal intensity of the MYCBP2 within the above fields of view. **(B-D)** The characteristics of MYCBP2 in boutons were analysed as follows: (B) MYCBP2 puncta area significantly altered by both α-Syn-WT and -A30P mutant overexpressions compared to control $F(2,32)=5.85$, $p=0.0068$; (C) MYCBP2 count within boutons did not significantly alter; (D) Relative intensity of MYCBP2 puncta was significantly reduced in the α-Syn-WT and -A30P conditions $F(2,34)=6.00$, $p=0.0057$. **(E-G)** The characteristics of MYCBP2 in axon segments were similarly analysed for (E) area size, (F) count and (G) intensity, however, no significant differences were observed in these conditions. **(H-J)** Two-way ANOVA analysis was conducted to examine interaction effects within the following conditions: (H) There was a significant interaction between the region measured and the puncta area $F(2,61)=5.09$, $p=0.0091$; (I) Significant interaction was not identified between the region measured and the puncta count; (J) The puncta intensity did not show a significant interaction with the region measured. Box plots: dashed lines indicate mean; colours indicate biological repeats; diamonds indicate data outside 1.5x IQR. Scale bar: 1 µm. Error bars: (H-J) SD. N=2-4 boutons from 32-37 cells. Images acquired on iSIM.

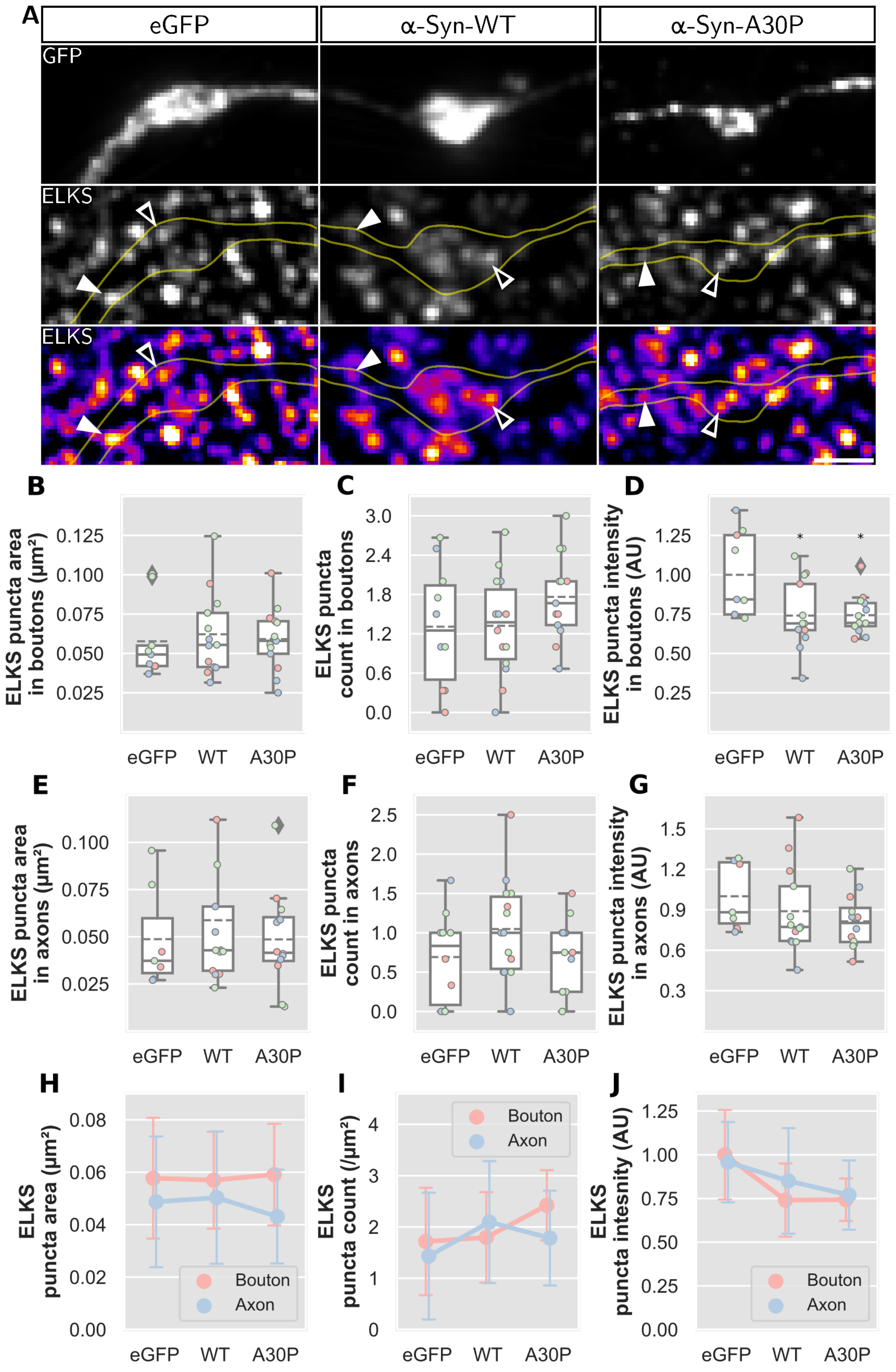

A
eGFP
α-Syn-WT
α-Syn-A30P
GFP
ELKS
ELKS
B
ELKS puncta area in boutons (μm²)
C
ELKS puncta count in boutons
D
ELKS puncta intensity in boutons (AU)
E
ELKS puncta area in axons (μm²)
F
ELKS puncta count in axons
G
ELKS puncta intensity in axons (AU)
H
ELKS puncta area (μm²)
I
ELKS puncta count (/μm²)
J
ELKS puncta intesnity (AU)
Bouton
Axon
eGFP
WT
A30P

**Figure 9 Effect of α-Synuclein overexpression of the organisation of the active zone marker ELKS. (A)** Rat cortical neurons were transfected with either eGFP, α-Syn-WT or -A30P. The cells were immunostained for both GFP and ELKS. Hollow arrows indicate ELKS lying within the bouton regions. The solid arrows show ELKS puncta distributed within the axon region. The bottom pseudo-coloured panels of ELKS demonstrate the signal intensity of the ELKS within the above fields of view. **(B-D)** The characteristics of ELKS in boutons between overexpression conditions were analysed as follows: (B) ELKS puncta area showed no significant alterations; (C) ELKS puncta count within boutons also showed no significant change; (D) The relative intensity of ELKS in the bouton was significantly altered in the α-Syn-WT and -A30P conditions $F(2,32)=5.28$, $p=0.0104$. **(E-G)** The characteristics of ELKS in axon segments were similarly analysed for (E) area size, (F) count and (G) intensity, however, no significant differences were observed in these conditions. **(H-J)** Two-way ANOVA analysis was conducted to examine interaction effects between the area analysed and the following conditions: (H) Puncta area; (I) Puncta count; (J) Puncta intensity. No significant interaction was observed in any of these conditions. Box plots: dashed lines indicate mean; colours indicate biological repeats; diamonds indicate data outside 1.5x IQR. Scale bar: 1 µm. Error bars: (H-J) SD. N=2-4 boutons from 35-37 cells. Images acquired on iSIM.

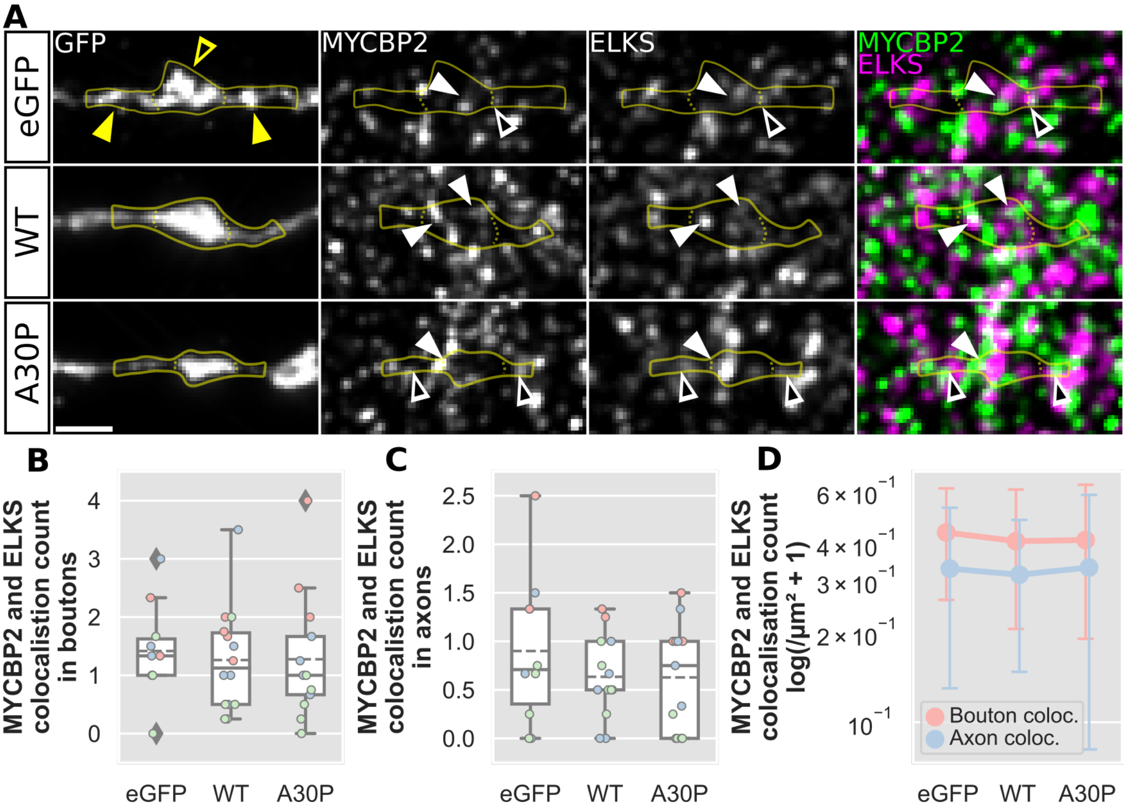

A
GFP
MYCBP2
ELKS
MYCBP2
ELKS
eGFP
WT
A30P
B
MYCBP2 and ELKS colocalistion count in boutons
C
MYCBP2 and ELKS colocalistion count in axons
D
MYCBP2 and ELKS colocalisation count log(/µm² + 1)
Bouton coloc.
Axon coloc.
eGFP
WT
A30P

**Figure 10 Colocalisation of MYCBP2 and ELKS and presynaptic sites. (A)** Rat cortical neurons were transfected with either eGFP, α-Syn-WT or -A30P overexpression plasmids. The regions that were measured are identified in the GFP panel with yellow arrows. The hollow yellow arrows indicate the bouton regions analysed, and the solid yellow arrows demonstrate the axon regions analysed. The last pseudo-coloured panels show MYCBP2 immunostaining in green and ELKS immunostaining in magenta. The overlap of these colours shows as white indicating sites of colocalisation. The solid arrows indicate colocalisation within the bouton regions, whereas the hollow arrows identify colocalised puncta in the axon regions. **(B-C)** There was no significant difference to the colocalisation count of MYCBP2 and ELKS in neither the boutons (B) or the axon regions (C). **(D)** There was no significant interaction between the region analysed and the type of overexpression on the colocalisation count of MYCBP2 and ELKS. Box plots: dashed lines indicate mean; colours indicate biological repeats; diamonds indicate data outside 1.5x IQR. Scale bar: 1 µm. Error bars: (D) SD. N=2-4 boutons from 36-37 cells. Images acquired on iSIM.
